# A switch in clathrin turnover controls endocytic coat size and organisation

**DOI:** 10.64898/2026.08.10.743920

**Authors:** Anne-Laure Boinet, Mamta, Anne-Sophie Rivier-Cordey, Aurélien Roux, Marko Kaksonen

## Abstract

Endocytosis internalises nutrients, regulates extracellular signals, and recycles membrane components. Clathrin polymerises into a coat that shapes the endocytic vesicle from the plasma membrane. However, the role of clathrin’s dynamic assembly in the endocytic process remains unclear. We show, using two-colour fluorescence recovery after photobleaching assays in yeast, that the clathrin coat turns over rapidly in the early phase of endocytosis, dependent on the auxilin Swa2 and its ATPase. In the late phase the turnover is stopped by the coat protein Sla1. Regulated clathrin turnover is critical for the timing of endocytic progression and for controlling coat size. In the absence of this dynamic regulation the endocytic coats become abnormally large, resulting in the failure of the final actin-driven vesicle budding. These findings reveal that, in addition to its classic structural function, the dynamic properties of the clathrin lattice are critical for both the temporal and mechanical aspects of endocytosis.

## INTRODUCTION

Clathrin-mediated endocytosis (CME) is a major and conserved internalisation route used by eukaryotic cells for the uptake of nutrients and signalling molecules, as well as for the recycling of membrane proteins. This process involves more than sixty different proteins that coordinate to form a coat that drives plasma membrane deformation and cargo selection to form a cargo-filled vesicle. Clathrin is the major component of this endocytic coat. Clathrin has been extensively studied since its initial discovery by Barbara Pearse in 1975 (Pearse, 1975), and its structure is well characterised: it is a triskelion-shaped heterohexameric protein that can assemble into large-scale polyhedral structures with variable curvature from flat lattices to fully formed spherical cages. Its properties allowing for its assembly into shapes of varying curvatures, as well as the timing of clathrin assembly at endocytic sites have been extensively studied, as well as its interactions with other endocytic proteins. Despite this large body of work over forty years (Kaksonen and Roux, 2018), the essential mechanical link between the biochemistry of clathrin and the generation of membrane curvature in cells remains to be identified: both *in vitro* and *in vivo*, curved and coated membrane structures arguing for a direct role of clathrin in membrane deformation, as well as flat membrane lattices arguing against, have been observed, and debated.

Clathrin is a rigid protein that can form flat hexagons, but also heptagons and pentagons associated with curved lattices. Therefore, how it assembles is intricately linked with the overall shape of the structure, as the position and the distribution of hexagons, pentagons and heptagons is topologically linked to the shape of the lattice. Understanding if clathrin changes the shape of membranes or stabilises the shape of membranes acquired by other means depends on how it is dynamically assembled.

Importantly, clathrin deletion impacts timing and robustness of membrane deformation in yeast, but does not completely abolish endocytosis (Kaksonen et al., 2005; Kukulski et al., 2016). Similar results are seen in mammalian cells (Hinrichsen et al., 2003), supporting the notion that clathrin does not directly participate in membrane bending, and that clathrin being a direct driver of membrane deformation might be too simplistic. Many studies focusing on clathrin dynamics at endocytic sites hint at a regulatory role for clathrin. In mammalian cells, clathrin turnover at endocytic sites was shown to be driven by the activity of its uncoating machinery, composed of a DnaJ protein, auxilin1/2, and an ATPase of the heat-shock cognate 71-kDa (Hsc70) protein family (He et al., 2020; He et al., 2025). Cell-free reconstitution experiments have demonstrated that impeding clathrin turnover at endocytic sites does not prevent pit formation, but strongly reduced transferrin receptor (TfR) clustering at those sites, suggesting a role in cargo recruitment (Chen et al., 2019). Recent studies in live cells showed that abolishing DnaJ/ATPase-driven clathrin turnover increases the proportion of short-lived abortive events and hinders pit formation (He et al., 2025; Krishnan et al., 2025). Taken together, these results highlight a link between clathrin coat remodelling and endocytic site maturation, that might not involve a direct membrane deformation activity.

In mammalian cells, pit formation can be variable both in its onset, as well as its duration (Sochacki and Taraska, 2019; Chen and Schmidt, 2020). Pit formation also depends on the cell type as well as growth conditions (Sochacki et al., 2021), making it a complex mechanism to study. In *Saccharomyces cerevisiae* however, it is easier to follow membrane deformation, that was shown to be highly stereotypical, consistently occurring during the last ∼10 seconds of the whole process (Kukulski et al., 2012). Moreover, clathrin was also shown to display a dynamic behaviour at endocytic sites in yeast cells (Newpher et al., 2005), making *S. cerevisiae* an ideal model to explore the interplay between clathrin assembly dynamics and CCP formation.

In this study, we sought to investigate clathrin dynamics at single endocytic sites throughout endocytic coat formation, and correlate clathrin dynamics with endocytic maturation and membrane deformation. We found that clathrin dynamics, in contrary to other endocytic proteins, transition from a high turnover to a solid state prior to the onset of the budding phase of endocytosis. By mutating endocytic factors involved in clathrin dynamics, in particular the DnaJ/Hsp70 uncoating machinery and late-coat protein Sla1, we further show that coat sizes are largely increased and budding is severely impaired. Altogether, our results support a model in which solidification of the clathrin lattice through controlled polymerisation allows for membrane deformation by actin forces. Overall, it shows that transitions from flexible to solid states are essential regulators of coats involved in membrane remodelling.

## RESULTS

### Clathrin displays a biphasic behaviour during endocytic site formation

We first sought to shed light on clathrin dynamics throughout the entire endocytic coat assembly process. We generated a *S. cerevisiae* strain expressing endogenously tagged clathrin light chain Clc1-GFP. Because clathrin is also involved in Trans-Golgi network (TGN) trafficking, we imaged clathrin at endocytic sites using total internal reflection fluorescence (TIRF) microscopy (FIG 1A). We photobleached Clc1-GFP at endocytic sites, and studied its fluorescence recovery after photobleaching (FRAP). We found two distinct outcomes: some photobleaching events were quickly followed by an increase in GFP signal, whereas others did not show any GFP signal recovery (FIG 1B). We hypothesised that these two behaviours might be correlated with the timeline of endocytic coat assembly; particularly, the events associated with an absence of Clc1-GFP signal recovery could be linked to the coat internalisation step, where the clathrin signal disappears from the evanescence field. In order to test that hypothesis, we imaged clathrin together with Ede1-mCherry or Pan1-mCherry as temporal markers, to visualise clathrin behaviour during the early and late phases, respectively (**FIG S1A**). Notably, since Pan1 assembly curves typically show a continuous increase until coat internalisation (Sun et al., 2015; Tolsma et al., 2018), a recovery of the Pan1-mCherry signal would indicate a photobleaching event occurring prior to budding. We compared the average FRAP for Clc1-GFP during the early and late phases, and found a striking difference between the two. Photobleaching events occurring during the early phase led to a quick recovery of ∼45% of the Clc1-GFP signal, with a half-time (t_1/2_) of ∼1.7 seconds (calculated from exponential fit). On the other hand, photobleaching events occurring during the late phase, and prior to the end of the endocytic coat assembly process (as shown by a recovery of the Pan1-mCherry fluorescence signal (**FIG S1B**)) were associated with an absence of fluorescence recovery for clathrin (FIG 1C). These results suggested that clathrin turns over during the early phase and becomes stable during the late phase.

**FIGURE 1:**
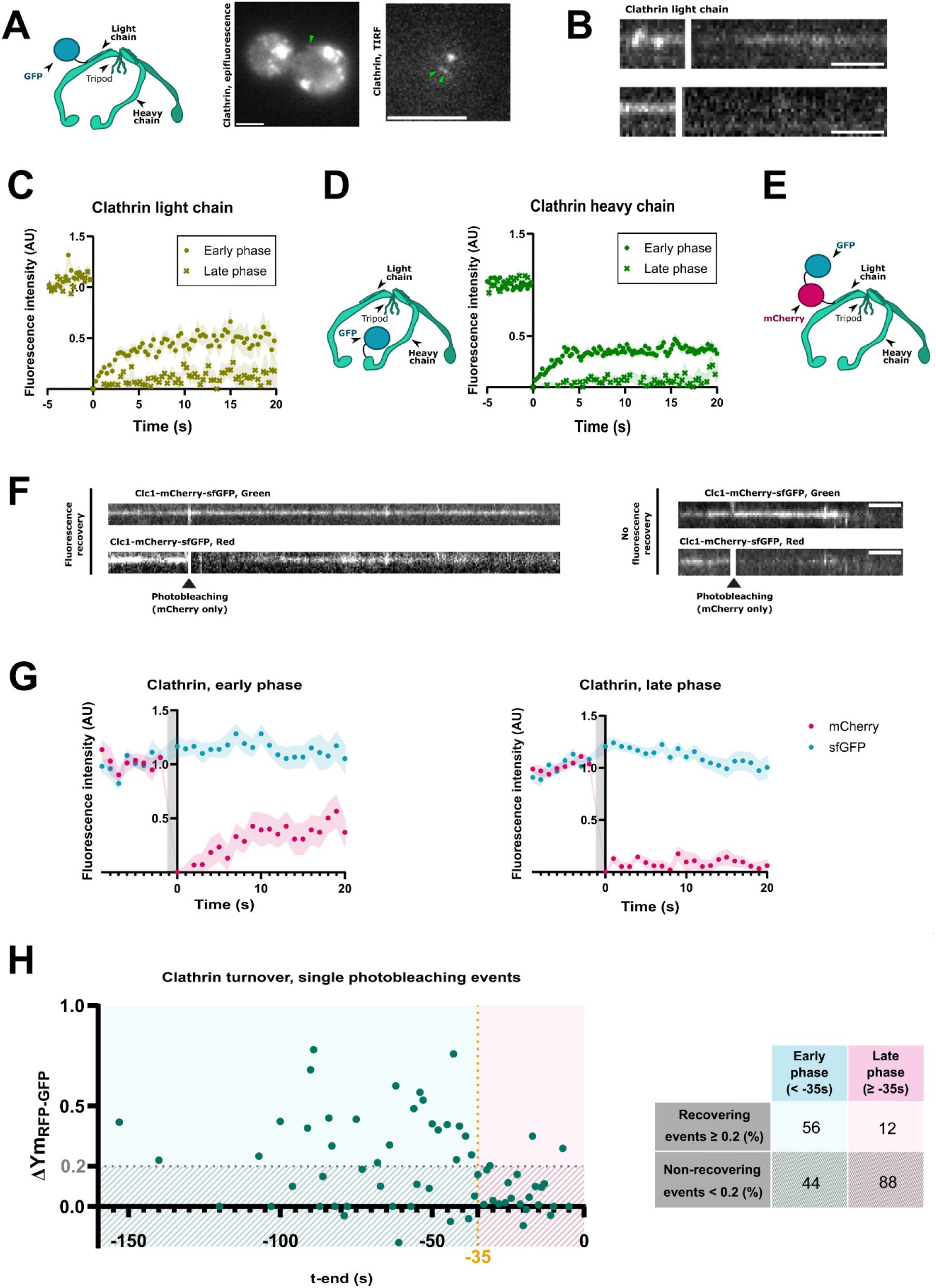
Clathrin dynamics at endocytic sites are biphasic. **A.** Left panel: schematic representation of the structure of a clathrin triskelion containing clathrin light chain (Clc1) tagged with GFP on its carboxyl end. Only one tagged clathrin light chain is represented for clarity. Right panels: Representative images of a *S. cerevisiae* cell expressing Clc1- GFP, acquired with widefield epifluorescence imaging of its equatorial plane, and with total internal reflection fluorescence microscopy (TIRF). Green arrows represent clathrin at endocytic sites. Scale bar: 4µm. **B.** Representative kymographs of Clc1-GFP fluorescence recovery after photobleaching (FRAP) events. Scale bar: 5 seconds. **C.** Mean Clc1-GFP FRAP (± standard error associated to the mean (SEM)) during the early phase (e.P.) compared to the late phase (l.P.). e.P.: N=64; l.P.: N=60. **D.** Left panel: schematic representation of the structure of a clathrin triskelion containing clathrin heavy chain (Chc1) tagged with GFP on its N- end. Only one tagged clathrin light chain is represented for clarity. Right panel: Mean GFP-Chc1 FRAP (±SEM) during the early phase compared to the late phase. e.P.: N=69; l.P.: N=78. **C,D.** Ede1-mCherry and Pan1-mCherry were used as early and late phase markers, respectively. **E.** Schematic representation of the structure of a clathrin triskelion containing Clc1 tagged with GFP and mCherry. Only one tagged clathrin light chain is represented for clarity. GFP allows for the tracking of protein amounts, while photobleaching mCherry provides information about clathrin turnover. **F.** Representative kymographs of Clc1-mCherry-sfGFP FRAP. Scale bar: 5 seconds. **G.** Mean mCherry FRAP (±SEM) of events occurring during the early phase (left; N=19) or the late phase (right; N=33), compared to mean GFP fluorescence intensity (±SEM), for clathrin at endocytic sites. **H.** Left: temporal distribution of mobile fractions from single photobleaching events, calculated from exponential fitting of single FRAP curves. T=0 represents the disappearance of the GFP signal, marking the end of the endocytic event. Right: proportion of clathrin photobleaching events leading to signal recovery or lack thereof, in the early phase v. the late phase.

Previous studies have shown distinct turnover properties for clathrin light and heavy chains in mammalian cells (Loerke et al., 2005). Therefore, we next wanted to determine whether the behaviour we observed during yeast endocytosis was that of the clathrin light chain specifically, or the whole clathrin triskelion. We generated a strain expressing endogenously tagged GFP-Chc1 (FIG 1D), and followed its FRAP during both early and late phases, using Ede1-mCherry and Pan1-mCherry as phase markers. We found a similar behaviour for the clathrin heavy chain to that previously observed for the light chain: during the early phase, photobleaching GFP-Chc1 was followed by a ∼37% signal recovery and a t_1/2_ of 1.4 seconds, while late-phase photobleaching events were associated with a lack of fluorescence recovery (FIG 1D). This demonstrated that clathrin triskelia turn over during the early phase of endocytosis, whereas the clathrin lattice is stable during the late phase.

Importantly, single-colour FRAP experiments, while being often used to determine protein dynamics, present one important limit when used for assembling proteins: separating turnover and growth is difficult. In our experiments, it remained hard to distinguish a stable assembly of the clathrin coat (i.e. clathrin triskelia being added progressively to the coat during the early phase, and the coat reaching its final size during the late phase) from a true biphasic behaviour. To study dynamics of clathrin throughout coat assembly, we optimised a simultaneous 2-colour FRAP experimental setup, in which the protein of interest is fused with mCherry and sfGFP (referred to as GFP) in tandem (**FIG 1E**). Using the two-colour labelling and photobleaching only mCherry, we can separate growth (variation in GFP signal) from turnover (difference in variation between mCherry and GFP). We first verified that using a 561nm laser photobleached mCherry, but not GFP (**FIG 1F**). We also verified that no Förster resonance energy transfer (FRET) due to acceptor photobleaching was detectable. We validated our assay by assessing the variability between GFP and mCherry fluorescence intensities of unbleached endocytic events for all endocytic proteins studied in this work. This revealed a high correlation between GFP and mCherry signal intensities over time (**FIG S1C, S2A**).

We endogenously tagged clathrin light chain Clc1 with mCherry and sfGFP in tandem in *S. cerevisiae* (**FIG 1E**). The tagging did not show significant impact on cell growth, clathrin localisation or endocytic progression, thus validating the functionality of the fusion protein (**FIG S1C, S1D**). We photobleached mCherry only and compared its fluorescence recovery to the change in GFP fluorescence intensity (**FIG 1F**). We separated the early-phase photobleaching events from the late-phase photobleaching events, and averaged their FRAP (**FIG 1G**). Like all early-arriving endocytic proteins, clathrin lifetimes vary from as little as 30 seconds to several minutes. However, clathrin intensity abruptly decreases at the final stage of vesicle budding as measured by TIRF microscopy because of the internalisation of the vesicle, setting the final point of endocytosis (**FIG S1C**). The late phase of the endocytic process consistently lasts 30 to 40 seconds, therefore we set the early-to-late phase transition point at 35 seconds before GFP signal disappearance, and assessed average clathrin dynamic properties during the two phases (**FIG 1G**). In both cases, the average GFP signal appeared largely constant. The average mCherry fluorescence intensity increased rapidly after photobleaching during the early phase. Again, this fluorescence recovery was partial, as shown by a calculated mobile fraction Ym ≈ 0.45, suggesting the coexistence of a dynamic clathrin and a stable clathrin populations within a single endocytic site. The average mCherry fluorescence intensity after photobleaching during the late phase, however, remained constant and close to 0.

To determine whether the difference in clathrin dynamics observed between the early and late phases results from a gradual change, or a switch in behaviour occurring at the transition time between the early and late phases, we aligned all photobleaching events using the GFP disappearance point as time zero, maintaining the early-to-late phase transition point at 35 seconds before GFP signal disappearance (**FIG 1H**). We plotted on a time axis the mobile fractions Ym determined from each single photobleaching event. Quite strikingly, photobleaching events leading to a significant fluorescence recovery (i.e. Ym > 0.2) almost exclusively occurred during the early phase, whereas those associated with a lack of signal recovery (Ym ≤ 0.2) primarily occurred during the late phase (**FIG 1H**). This indicates that the switch in dynamic behaviour overlapped with the time of early-to-late phase transition.

Altogether, our results indicate that the clathrin lattice switches from a dynamic structure to a stable one at the transition between the early and late phases, meaning that clathrin coat stabilisation precedes the onset of membrane deformation.

### Endocytic site maturation is associated with an increased coat stability

We next sought to verify whether the biphasic dynamic behaviour is specific to clathrin, or is common to other endocytic proteins. We used our 2-colour FRAP assay to study the turnover of endocytic proteins from various functional modules, assessing general dynamic properties of the endocytic coat. We endogenously two-colour labelled early protein Ede1, mid-coat protein Sla2, late-coat protein Sla1, and actin-module protein Abp1 (**FIG 2**). All proteins displayed varying turnover rates, with Ede1 and Abp1 appearing to be the most dynamic, and Sla1 the most stable. None of the proteins studied changed their behaviour throughout the course of endocytic site maturation (**FIG 2A-D**). Although each endocytic protein appeared to be characterised by distinct turnover parameters, we could still draw general conclusions about the dynamics of the maturing endocytic assembly. Proteins from the early coat, Ede1 (**FIG 2A**) and, as shown previously, clathrin during the early phase (**FIG 1**), displayed high turnover rates and high mobile fractions. On the contrary, mid-coat protein Sla2 (**FIG 2B**), as well as late-coat protein Sla1 (**FIG 2C**), displayed little to no turnover. Finally, actin crosslinker Abp1 (**FIG 2D**), was associated with a fast protein turnover. Additional 2-colour FRAP experiments on early-coat clathrin adaptor Yap1801, mid-coat protein Ent1, late-coat proteins Pan1 and Las17 as well as actin motor Myo5, revealed similar dynamic behaviours to those observed for proteins from the same functional modules (**FIG S2B**).

**FIGURE 2:**
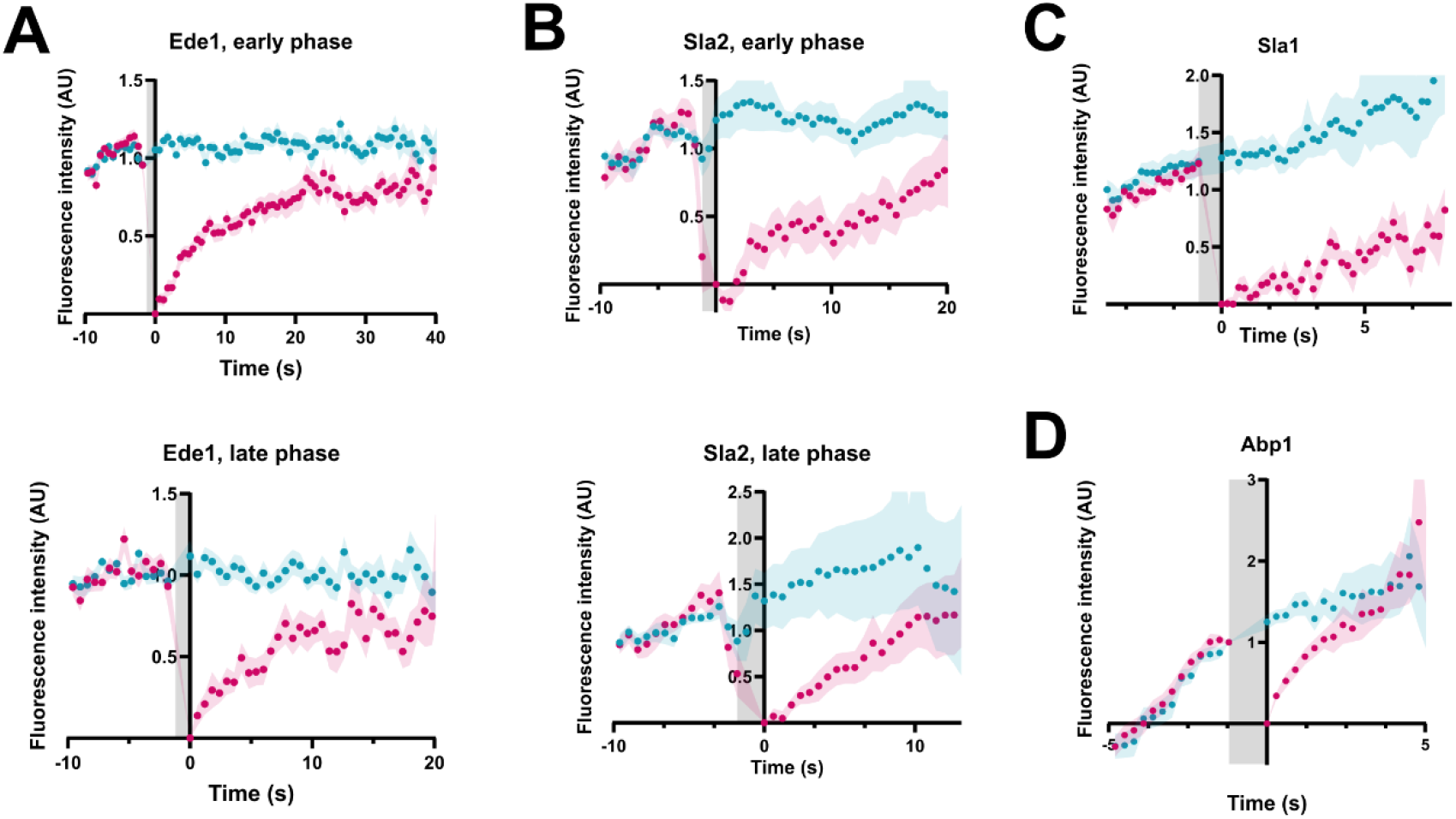
Endocytic coat maturation correlates with its increased stability. A-D. Average mCherry FRAP compared to GFP fluorescence intensity. Gray areas represent photobleaching durations. Error bars represent standard errors associated with the mean (SEM). For early protein Ede1 (**A**; N(e.P.)=52; N(l.P.)=18), and mid-coat protein Sla2 (**B**; N(e.P.)=21; N(l.P.)=41), photobleaching events occurring during early and late phases were separated and represented on distinct plots. All photobleaching events for late coat protein Sla1 (**C**; N=18) and Actin-module protein Abp1 (**D**; N=51) were represented on single plots.

Taken together, these results suggest that the various endocytic modules that were initially characterised by their recruitment time (Kaksonen et al., 2005), have specific dynamic properties as well. Proteins from the early phase display high turnover rates, while in the late phase, the coat associated proteins are largely stable. Proteins associated with the actin network that pulls on the coat are highly dynamic.

### The Swa2/ATPase uncoating machinery drives clathrin turnover during the early phase

We next sought to further characterise the mechanism of clathrin turnover during the early phase. In yeast, auxilin homolog Swa2 was shown to play a crucial role in clathrin coat disassembly (Gall et al., 2000). We generated a mutant strain expressing Swa2 lacking its J domain (*swa2-ΔJ*), and therefore unable to recruit the ATPase to clathrin structures (Gall et al., 2000; Xiao et al., 2006; FIG 3A**, S3**). Epifluorescence imaging of clathrin-GFP in the *swa2-ΔJ* background highlighted the presence of many small, mobile puncta throughout the cytosol, corresponding to clathrin-coated vesicles. This phenotype, which was also observed upon Swa2 deletion, suggests a wide clathrin uncoating defect (Gall et al., 2000; **FIG S3B**).

**FIGURE 3:**
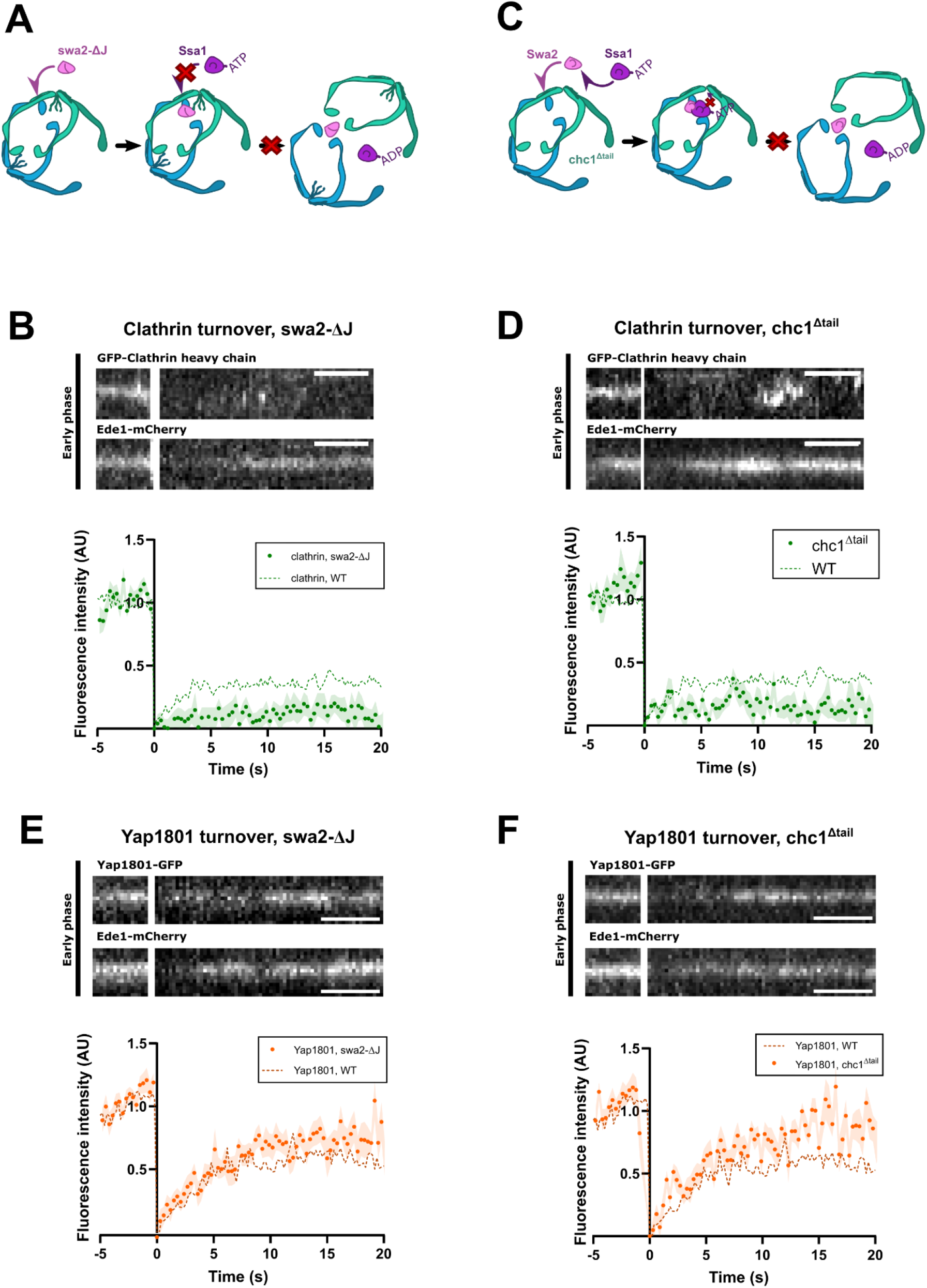
Early-phase clathrin turnover is driven by the Swa2/ATPase uncoating machinery. **A.** Schematic represantation of clathrin uncoating hampered by swa2-ΔJ mutation. Swa2 lacks its J-domain, and is therefore unable to recruit the uncoating ATPase to clathrin. **B.** Clathrin heavy chain FRAP during the early phase, in cells expressing swa2-ΔJ. Top: representative kymograph of a photobleaching event. Bottom: mean FRAP (±SEM) of N=32 events (swa2-ΔJ) compared to WT (N=64). Ede1-mCherry is used as phase marker. **C.** Schematic representation of clathrin uncoating hampered by chc1^Δtail^ mutation. Swa2 binds to clathrin and recruits the uncoating ATPase, but the ATPase cannot bind to the missing C-term unstructured tail. **D.** Clathrin FRAP during the early phase, in cells expressing chc1^Δtail^. Top: representative kymograph of a photobleaching event. Bottom: mean FRAP (±SEM) of N=18 events (chc1^Δtail^) compared to WT (N=64). Ede1-mCherry is used as phase marker. **E,F.** Yap1801 FRAP during the early phase, in cells expressing swa2-ΔJ (**E**) or chc1^Δtail^ (**F**). Top: representative kymograph of a photobleaching event. Bottom: mean Yap1801 FRAP (±SEM) in cells expressing swa2-ΔJ (E, N=40) or chc1^Δtail^ (F, N=24) compared to WT (N=43). Ede1-mCherry is used as phase marker.

We observed that the clathrin-GFP turnover was strongly reduced during the early phase in the *swa2-ΔJ* background (FIG 3B). We also generated a yeast strain expressing a truncated clathrin heavy chain chc1^Δtail^, lacking the unstructured tail in its carboxyl end that was shown to bind the ATPase Hsc70 and stimulate its uncoating activity in mammals (FIG 3C**, S3**) (Rapoport et al., 2007). Imaging of clathrin-GFP in the chc1^Δtail^ background revealed a very similar phenotype to the swa2-ΔJ mutation, with a high number of small, mobile puncta throughout the cytosol (**FIG S3B**). Clathrin turnover was also strongly reduced in the chc1^Δtail^ mutant, confirming the conservation of the binding mechanism between clathrin and the ATPase (**FIG 3D**). We also observed a strong reduction of clathrin turnover upon ATP depletion, further supporting the involvement of the uncoating machinery in the early phase clathrin turnover (**FIG S3C**).

We next wanted to know how preventing clathrin dynamics during the early phase would affect the turnover of clathrin adaptors. We investigated the impact of clathrin turnover inhibition on adaptor protein Yap1801. Strikingly, neither *swa2-ΔJ* (**FIG 3E**) nor chc1^Δtail^ (**FIG 3F**) had any effect on Yap1801 turnover properties, suggesting that clathrin adaptor dynamics are independent from clathrin dynamics.

Altogether, these results show that the clathrin disassembly machinery Swa2/DnaJ is required for the turnover of clathrin during the early phase of endocytosis.

### Sla1 stabilises clathrin upon late-phase transition

We next investigated the mechanism behind clathrin’s stabilisation upon late phase transition. Interactions with a partner recruited at the onset of the late phase could explain this stabilisation. Upon transition to the late phase, the first proteins to be recruited to the endocytic site are Pan1, Sla1, End3 and Las17 (Kaksonen et al., 2005; Sun et al., 2015). Out of these proteins, only Sla1 interacts with clathrin, via an LLDLQ clathrin-binding motif (CBM) (Di Pietro et al., 2010; **FIG S4A**). Additionally, in absence of Sla1, endocytosis can proceed, however the recruitment of the proteins from the actin module is delayed (Warren et al., 2002; Kaksonen et al., 2005).

In order to determine whether Sla1 is involved in late-phase clathrin coat stabilisation, we assessed clathrin dynamics in a *sla1Δ* background, using Pan1-mCherry as a late phase marker (FIG 4A). We observed that in absence of Sla1, clathrin continuously turned over throughout the late phase, and with dynamic properties similar to the early phase in wild type cells. This indicates that when Sla1 is not present at endocytic sites, the clathrin coat fails to stabilise.

**FIGURE 4:**
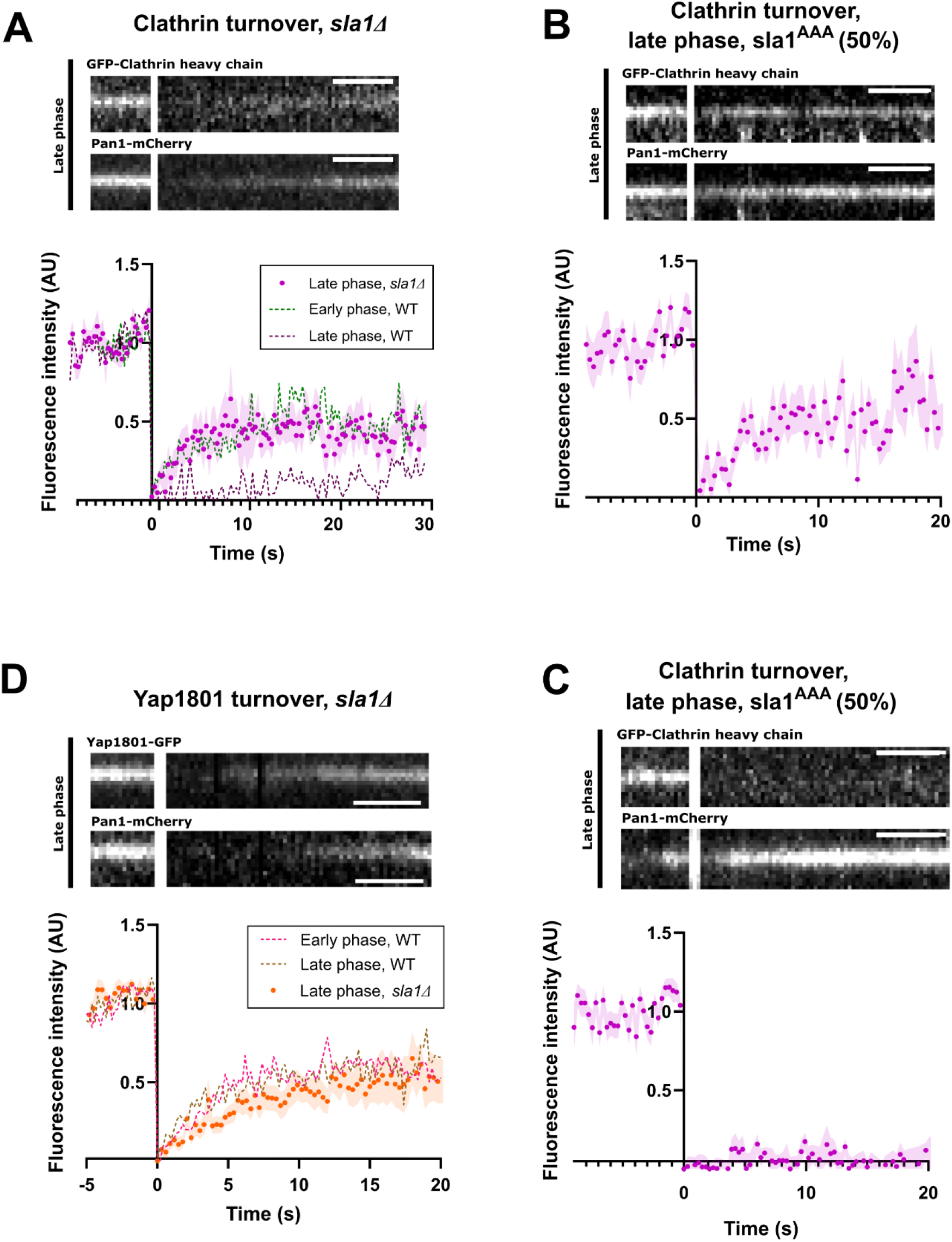
Sla1 stabilises the clathrin coat during the late phase. **A.** Clathrin heavy chain FRAP during the late phase, in absence of Sla1. Top: representative kymograph of a photobleaching event. Bottom: mean FRAP (±SEM) of N=24 events (*sla1Δ*) compared to WT (e.P.: N=64; l.P.: N=60). Pan1-mCherry is used as phase marker. **B, C.** Clathrin FRAP during the late phase, in cells expressing sla1^AAA^. Top: representative kymograph of a photobleaching event. Bottom: mean clathrin FRAP (±SEM) in cells expressing sla1^AAA^ of events showing fluorescence recovery (**B**, N=29) or an absence of recovery (**C**, N=32). Pan1-mCherry is used as phase marker. **D.** Yap1801 FRAP during the late phase, in absence of Sla1. Top: representative kymograph of a photobleaching event. Bottom: mean FRAP (±SEM) of N=28 events (*sla1Δ*) compared to WT (e.P.:N=43; l.P.: N=35). Pan1- mCherry is used as phase marker.

To further test the role of Sla1-clathrin interaction, we generated a sla1^AAA^ mutant, harbouring a LLDLQ-to-AAALQ mutation at amino acid positions 803-805 in the Sla1 sequence, thus abolishing Sla1-to-clathrin binding (**FIG S4A**, Di Pietro et al., 2010). We characterised clathrin dynamics during the late phase in this background (FIG 4B**, C**). We observed a clear effect on clathrin FRAP in the sla1^AAA^ mutant: 48% of the endocytic events were characterised by clathrin dynamics similar to those observed in *sla1Δ* cells (**FIG 4B**). This suggests that Sla1-to-clathrin binding is directly involved in stabilising the clathrin coat. The other 52% of the endocytic events appeared normal, and were associated with a lack of clathrin turnover in the late phase (FIG 4C), which suggests that another mechanism exists to support Sla1-to-clathrin binding in stabilising the clathrin coat.

We next sought to verify whether changes in clathrin turnover during the late phase would impact that of its adaptor Yap1801. Both Sla1 deletion (**FIG 4D**) and sla1^AAA^ (**FIG S4B**) did not appear to affect Yap1801 dynamics during the late phase.

Taken together, these results show that Sla1 stabilises clathrin specifically via their interaction, without affecting the dynamics of clathrin adaptor Yap1801. This finding strengthens our conclusion that clathrin and its adaptors are part of two independent functional layers of the endocytic coat.

### Clathrin turnover during the early phase regulates endocytic site maturation

Since the turnover of clathrin depended on the disassembly machinery, we investigated the effects of swa2-ΔJ on the early and late phases of endocytosis, as well as the transition from early to late phase. We used Ede1-GFP and Pan1-mCherry as markers for the early and late phases, respectively (FIG 5A). In wild-type cells, endocytic site maturation ranges from ∼1-2 min, and consistently leads to vesicle internalisation and coat disassembly. In the *swa2-ΔJ* background however, the endocytic events were significantly more variable: while a portion was wild-type-like, we observed a large population of stalled and repeated events (FIG 5A**, B**).

**FIGURE 5:**
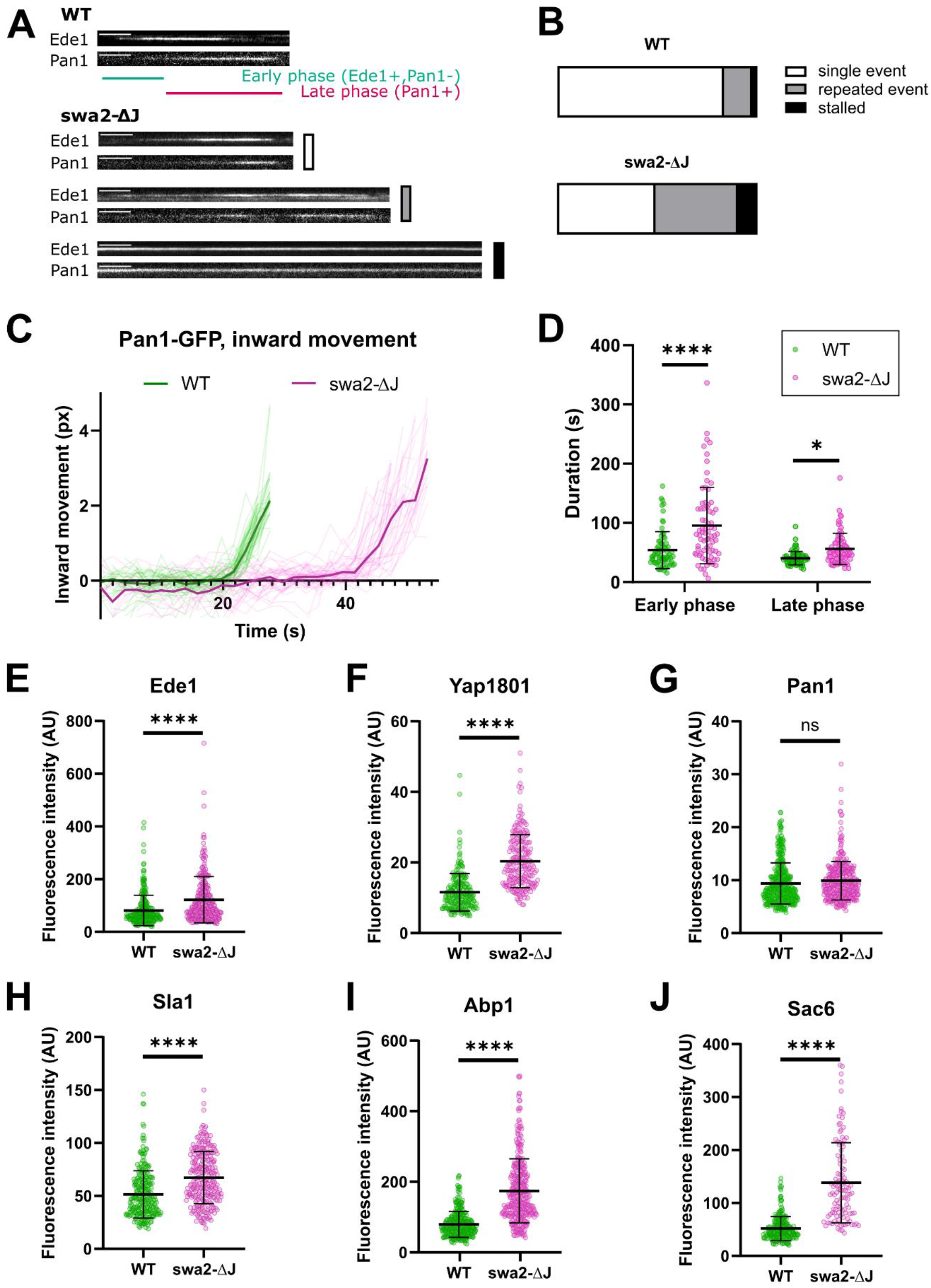
Impaired early-phase clathrin turnover hampers endocytic progression. **A.** Representative kymographs of early and late phase progression using Ede1-GFP and Pan1-mCherry as phase markers, in WT compared to swa2-ΔJ. Scale bars: 10 seconds. **B.** Quantification of the endocytic phenotypes highlighted in A. in WT cells compared to swa2-ΔJ mutants. WT: N=83, swa2-ΔJ: N=72. A,B. White: single events; gray: repeated events; black: stalled events. **C.** Pan1-GFP centroid tracking until coat internalisation, in WT compared to swa2-ΔJ. Thin curves represent individual tracks, bold curves represent the average of N=220 (WT) and N=218 (swa2-dJ) tracks. **D.** Quantification of early and late phase durations in WT compared swa2-ΔJ cells imaged with TIRF, using Ede1-GFP and Pan1-mCherry as phase markers. WT: N=75; swa2-ΔJ: N=75. P-values from Sidak’s test: early phase: <0.0001; late phase: 0.41. **E-J:** Mean fluorescence intensity of endocytic proteins Ede1 (**E**; N(WT/swa2-ΔJ)=282/266), Yap1801 (**F**; N(WT/swa2-ΔJ)=233), Pan1 (**G**; N(WT/swa2-ΔJ)=419/359), Sla1 (**H**; N(WT/swa2-ΔJ)=259/261), Abp1 (**I**; N(WT/swa2- ΔJ)=307/360) and Sac6 (**J**; N(WT/swa2-ΔJ)=246/111) in WT v. swa2-ΔJ cells. Fluorescence intensities are calculated from Z-projections of 15-seconds movies.

This defect in endocytic progression prompted us to first investigate the late internalization phase in the *swa2-ΔJ* mutant by centroid tracking of coat marker Pan1-GFP (FIG 5C). In the *swa2-ΔJ* background, although the average behaviour remained unchanged compared to wild-type, the time and speed of internalisation appeared to be much more variable. This suggests that early-phase clathrin turnover is an important player in the robustness of endocytosis, and in particular of the late internalisation phase. In support of this, the early phase was much longer on average when clathrin turnover was hampered, and its duration also was more variable (FIG 5D).

We also observed a defect in the early-to-late phase transition itself. In wild type cells, transition from early to late phase is associated with the recruitment of Pan1, Sla1 and End3, and quickly followed by the complete disassembly of Ede1. In the *swa2-ΔJ* background, Ede1 failed to disassemble in many events, and was still present at the time of budding (**FIG 5A, S5**). This Ede1 disassembly defect was also associated with higher protein amounts of early coat proteins, such as Ede1 and Yap1801 (**FIG 5E, F**), as well as late phase proteins, including Sla1 (**FIG 5H**) and, to an even stronger extent, actin-associated proteins Abp1 and Sac6 (**FIG 5I, J**). Of all proteins assessed, only Pan1 recruitment was unaffected (FIG 5G). Taken together, these results suggest that early-phase clathrin turnover is critical for the timing of the transition from early to late phase, potentially by regulating the amount of late proteins being recruited to the endocytic site. This regulation of endocytic progression and protein assembly, particularly those associated with actin assembly, appears to play a crucial role in the robustness of coat internalisation.

### Late-phase clathrin stabilisation regulates actin network assembly and vesicle budding

To understand the function of late-phase clathrin stabilisation, we investigated whether impeding it would affect membrane deformation. We chose to study the effects of sla1^AAA^ on endocytic progression; although the effects of this point mutation relating to clathrin stabilisation were milder, the phenotypes associated with this mutation would be more specific to its interaction with clathrin.

We quantified the proportion of endocytic events leading to successful coat internalisation in a yeast strain expressing sla1^AAA.^ In this strain, when compared to wild type, endocytic events were associated with a significantly longer late phase (FIG 6A**, C**). Moreover, a significant proportion of events were either stalled or repeated (**FIG 6B**), a phenotype reminiscent to that observed with the *swa2-ΔJ* mutation, albeit to a lesser extent. We next sought to determine whether this abnormal late phase was also associated with a defect in membrane deformation. We tracked the centroid of Pan1- EGFP in a strain harbouring the sla1^AAA^ mutation, and compared it to wild type (**FIG 6D**). Similar to what we had observed in the *swa2-ΔJ* mutant, we found an increased variability in the internalisation speed of the Pan1-EGFP patch, suggesting a defect in the robustness of membrane deformation.

**FIGURE 6:**
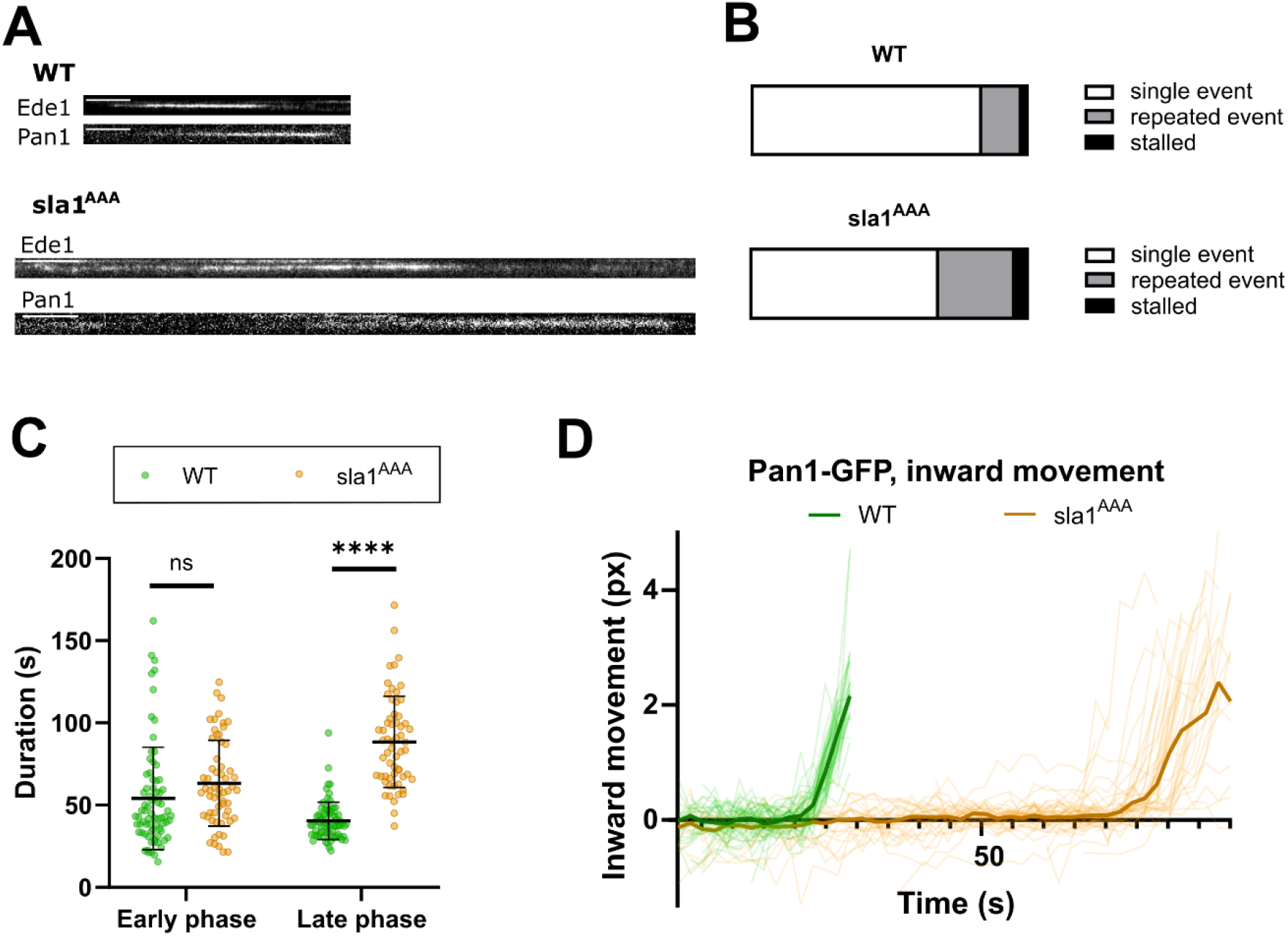
Impaired late-phase clathrin stabilisation hampers late-phase progression. **A.** Representative kymographs of early and late phase progression using Ede1-GFP and Pan1-mCherry as phase markers, in WT compared to sla1^AAA^. Scale bars: 10 seconds. **B.** Quantification of the endocytic phenotypes highlighted in A. in WT cells compared to sla1^AAA^ mutants. **C.** Quantification of early and late phase durations in WT compared sla1^AAA^ cells imaged with TIRF, using Ede1-GFP and Pan1-mCherry as phase markers. WT: N=75; sla1^AAA^: N=60. P-values from Sidak’s test: early phase: 0.9996; late phase: <0.0001. **D.** Pan1-GFP centroid tracking until coat internalisation, in WT compared to sla1^AAA^. Thin curves represent individual tracks, bold curves represent the average of N=51 (WT) and N=45 (sla1^AAA^) tracks.

Previous study on the impact of sla1^AAA^ mutant on endocytic coat formation revealed that late-coat proteins, including sla1^AAA^ itself, Pan1 and Las17, as well as actin-module proteins Abp1 and Myo5, were recruited at increased levels to endocytic sites (Tolsma et al., 2018).

Taken together, these results suggest that clathrin stabilisation that takes place at the beginning of the late phase regulates the assembly of the late-phase proteins to ensure effective vesicle budding.

### Controlled clathrin assembly provides a template for optimal endocytic site organisation

To understand the interplay between the two phases of clathrin assembly, we generated a double mutant defective for both early-phase clathrin turnover and late-phase clathrin stabilisation, and studied how endocytosis is affected. We abolished both Swa2/ATPase activity and Sla1-driven clathrin stabilisation by generating a *swa2-ΔJ sla1Δ* mutant strain, and observed a strong decrease in cell fitness compared to wild type *S. cerevisiae*: cell division was visibly hampered, leading to abnormal cell shapes (**FIG S6**). We focused on the impact of the double mutation on the endocytic process using late coat protein Pan1- GFP and actin crosslinker Sac6-mCherry as markers (FIG 7A). Both proteins appeared to be recruited to endocytic sites and subsequently internalised, although after a longer time compared to wild type, suggesting that endocytosis, was taking place (FIG 7A). Nevertheless, the fluorescence intensity of both Pan1 and Sac6 at endocytic sites had a striking increase and a broader distribution (**FIG 7A, B**) compared to wild-type cells (**FIG 7B**). Some of the endocytic patches were so large that we could resolve their extension along the membrane (FIG 7C), above the diffraction limit, therefore much larger than the typical 30-50nm of wild-type endocytic coats (Mund et al., 2018; Skruzny et al 2020). The timing, speed and depth of coat internalisation also appeared to be more variable than in wild type, to a much higher extent than in both single mutations (**FIG 7D**). Notably, the largest patches were associated with an aberrant internalisation behaviour (FIG 7C): the edges of the patches appeared to move inwards at a first, before the middle portion of the coat, giving rise to coats with concave side towards the cytoplasm, opposite of the normal curvature of endocytic coats. This prompted us to investigate the shape of the vesicle formed by such abnormal coat. We used correlative light and electron microscopy (CLEM) to visualise endocytic vesicles in the *swa2-ΔJ sla1Δ* mutant, using Abp1-GFP as a fluorescence marker. In wild type cells, the Abp1-GFP signal was associated with either endocytic pits or detached vesicles in >99% percent of events, both surrounded by a ribosome exclusion zone that corresponds to the actin network (Kukulski et al., 2012; **FIG 7E**). In the *swa2-ΔJ sla1Δ* mutant, the Abp1-GFP signal was associated with an assembled actin network, most of the time without any sign of membrane deformation (FIG 7F). The ribosome exclusion zones were also substantially larger than in wild type cells, suggesting larger actin networks – a result in line with the accumulation of Sac6 observed in fluorescence microscopy (**FIG 7A**). Comparing these results to the inward coat movement observed in live fluorescence microscopy, strongly suggests that in most cases, the coat becomes too large to deform the membrane and instead detaches from it by curling inward. These results indicate that controlling clathrin turnover is a crucial mechanism for the regulation of the size of endocytic coat and its ability to form vesicles.

**FIGURE 7:**
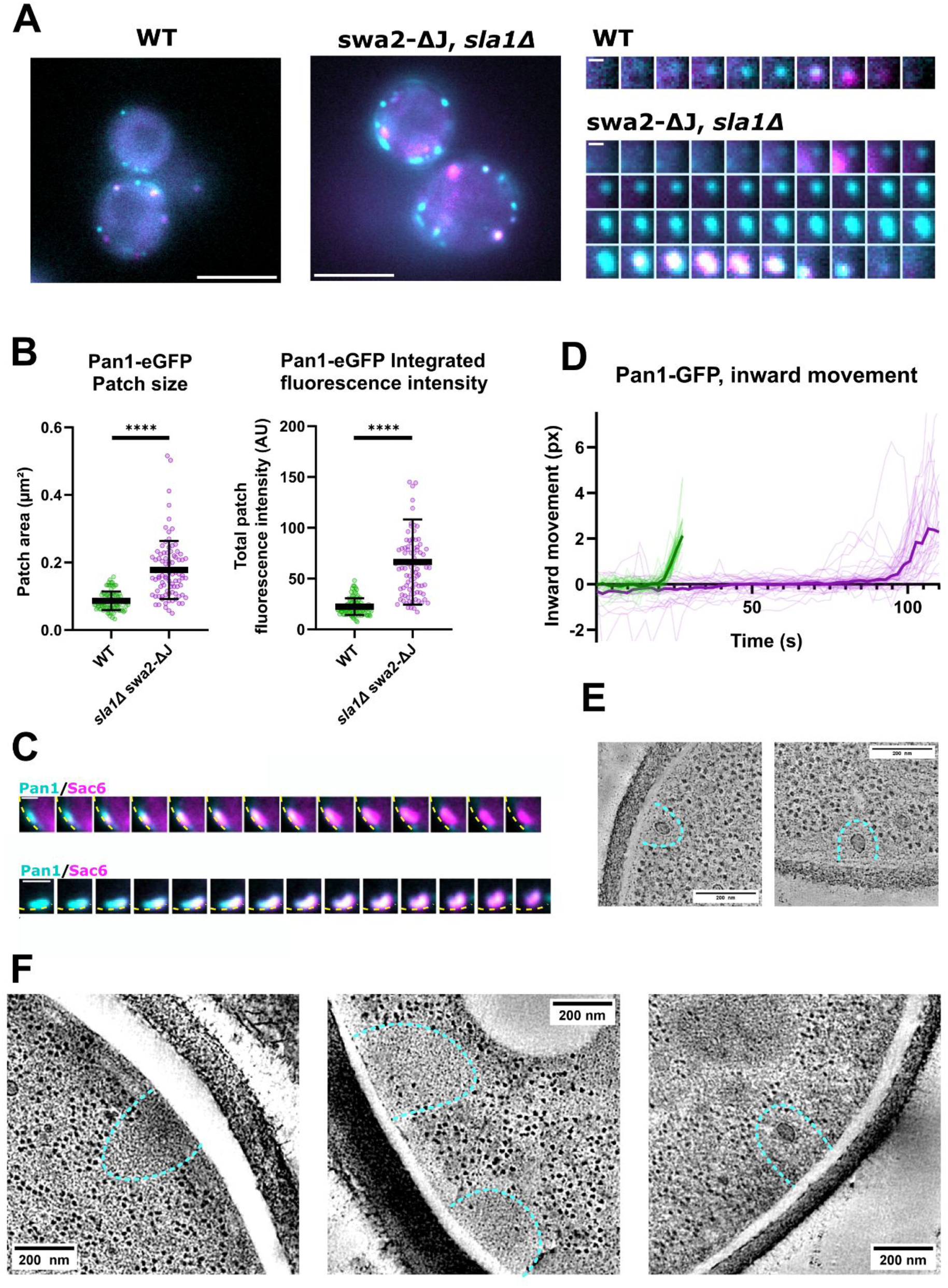
Uncontrolled clathrin assembly hampers endocytic vesicle formation. **A.** Representative images of WT (left) and *swa2-ΔJ sla1Δ* (middle) cells expressing Pan1-GFP (turquoise) and Sac6-mCherry (magenta). Brightness and contrast settings are adjusted independently for visibility. Scale bar: 4µm. Right: representative timelapse of endocytic events observed in WT (top) v. *swa2-ΔJ sla1Δ* (bottom) cells. Scale bar: 0.2µm. dt=7 seconds. **B.** Quantification of Pan1-GFP patch area (left; P-value from unpaired t-test with Welch’s correction <0,0001) and integrated fluorescence intensity (right; P-value from unpaired t-test with Welch’s correction <0,0001) in WT v. *swa2-ΔJ sla1Δ* cells. Measurements for single patches are represented, together with mean ± standard deviation (SD). WT: N=93; *swa2-ΔJ sla1Δ*: N=91. **C.** Two representative endocytic event timelapses observed in *swa2-ΔJ sla1Δ* cells, associated with endocytic coat growth beyond the size diffraction limit and apparent anisomorphic actin network assembly, demonstrated by Pan1-GFP and Sac6-mCherry, respectively. Yellow dotted lines highlight the cell contour. Total duration: 28 seconds. Scale bar: 0.2µm. **D.** Pan1-GFP centroid tracking until coat internalisation, in WT compared to *swa2-ΔJ sla1Δ*. Thin curves represent individual tracks, bold curves represent the average of N=51 (WT) and N=47 (*swa2-ΔJ sla1Δ*) tracks. **E.** Single frames from tomograms of endocytic sites from WT cells expressing Abp1-GFP, obtained by correlative light and electron microscopy (CLEM). Turquoise dotted lines highlight the ribosome exclusion zones. Scale bars: 0.2µm. **F.** Single frames from tomograms of endocytic sites from *swa2-ΔJ sla1Δ* cells expressing Abp1-GFP, obtained by CLEM. Turquoise dotted lines highlight the ribosome exclusion zones. Scale bars: 0.2µm.

## DISCUSSION

In this study, we have found that one of the prime structural proteins in endocytosis, clathrin, plays a direct role in the regulation of the organisation and size of the endocytic protein assembly through its dynamic properties. We have found that clathrin assembly dynamics are tightly controlled throughout endocytic coat formation. During the early phase, the clathrin coat is partially and rapidly turning over. This clathrin turnover depends on targeted removal of clathrin triskelia by the uncoating machinery Swa2/ATPase. Upon early-to-late phase transition, the clathrin coat is stabilised via the recruitment of late coat protein Sla1. Direct Sla1 binding to clathrin plays a crucial role in stabilising the clathrin coat. The regulation of clathrin dynamics controls the organisation and amount of coat proteins at the endocytic site and is therefore essential for controlled progression of endocytosis.

### Clathrin turnover regulates coat size

The mechanism defining the size of the clathrin coat and thus the resulting vesicle has remained unknown. A classic proposal for such mechanism is that the geometric closure of the clathrin coat that has a defined curvature determines the ultimate size of the coat. However, in yeast cells clathrin assembles on a flat membrane and the membrane shaping only happens during the last ten seconds of the endocytic process (Kukulski et al., 2012). In addition, in mammalian cells the strict constant curvature model has been challenged (Avinoam et al., 2015; Sochacki, et al. 2021). Coat assembly on the flat membrane would have no intrinsic limit for its size. We showed that the dynamic exchange of clathrin in the early phase and stabilisation of the clathrin lattice in the late phase are critical for proper size control of the coat. Interfering with either the early phase turnover or late phase stability leads to increased amounts of endocytic proteins to be assembled at the endocytic sites. In the double mutant where both early turnover and late stabilisation are prevented the endocytic coat continues to grow uncontrolled until actin assembly terminates the endocytic event. Rapid turnover of clathrin triskelia during the long early phase in an ATP-dependent manner means there is a significant energetic cost for endocytic events. Our findings suggest that this cost is required for the temporal and spatial control of endocytic protein assembly within the coat.

### The clathrin lattice templates the assembly of the endocytic machinery

Our results reveal a clear role for a regulated clathrin turnover in driving the recruitment and organisation of other endocytic proteins. Comparing the effects from clathrin turnover inhibition to those of a full clathrin deletion highlights the key roles played by clathrin in endocytic site formation. Absence of clathrin leads to a defect in initiation and proper maturation timing, as well as scission (Newpher et al. 2005; Kaksonen et al., 2005; Kukulski et al., 2016). Invagination speed and length are not affected and vesicle formation still occurs, albeit less consistently: the vesicles formed are more variable in size, which suggests a direct role for clathrin in the control of the size of the coat, and the vesicle it eventually forms. We have observed a similar phenotype in the absence of a controlled clathrin assembly: coat sizes are more variable and often highly enlarged compared to wild type. The stronger effect of uncontrolled clathrin assembly compared to its complete absence on coat organisation and vesicle formation, highlights that clathrin is a particularly potent director of endocytic coat assembly. The assembly of endocytic proteins appears to follow the organisation of the clathrin coat: when clathrin assembly is regulated, the assembly of the whole endocytic coat is efficient, and leads to the formation of a specific vesicle size (Kukulski et al., 2016). When clathrin is absent, other proteins tend to assemble into their preferred organisation, which is probably close to wild type; however their assembly is less efficient and less robust, which leads to changes in maturation time and coat/vesicle sizes. Strikingly, when clathrin is present, but its turnover is not regulated by Swa2/ATPase and Sla1, neither is the assembly of other endocytic proteins, thus hindering vesicle formation.

In addition to regulating the endocytic protein assembly, cargo selection may also depend on the dynamic clathrin coat. Cargo capture has been suggested to happen during the early phase (Carroll et al., 2012). Moreover, clathrin turnover was shown to drive cargo selection in cell-free reconstitution systems (Chen et al., 2019). Cargo binding might change the strength of clathrin-adaptor binding, which could in turn regulate Swa2/ATPase uncoating. Interestingly, the Swa2 sequence contains a ubiquitin-associated (UBA) domain that does not have any known function. Since ubiquitination is a well-established internalisation signal, a potential role for this UBA domain in clathrin triskelia targeting would be worth exploring.

Swa2’s role in the disassembly of the clathrin coat decorating endocytic vesicles both *in vitro* and in cells is well documented (Gall et al., 2000; Xiao et al., 2006; Krantz et al., 2013). We showed that Swa2-mediated mechanism drives the turnover of about half the clathrin lattice during the early phase. The simplest explanation for this partial clathrin turnover lies in the organisation of the coat: triskelia at the rim of the coat are probably more accessible than the ones at the centre and, because they have fewer interacting partners, their removal from the structure might be easier.

### Coat stability and actin force transmission

Using the two-colour FRAP that distinguished between turnover and growth, we found that endocytic proteins recruited during the early phase, namely Ede1 and Yap1801, are highly dynamic. The middle coat protein Sla2 also turned over, albeit to a lesser extent. On the other hand, middle coat protein Ent1, as well as late coat proteins Pan1, Sla1 and Las17 were stably recruited, and did not show any significant turnover. The stable assembly of late coat proteins around the dynamic early proteins, would therefore lead to an overall increase in endocytic coat stability. This coat stabilisation is followed by the recruitment of actin associated proteins, such as Abp1 and Myo5, which we found to be highly dynamic. Hampering late-phase clathrin coat stabilisation leads to defects in actin module assembly and coat internalisation. This phenotype points to a more general role for endocytic coat stabilisation in directing and supporting actin network polymerisation for membrane deformation.

In yeast cells, the actin network polymerisation provides most of the force needed to counteract turgor pressure (Dmitrieff et al., 2015). Although absence of clathrin turnover negatively impacts actin network organisation, actin assembly does not depend on clathrin, as vesicles can be formed even in absence of clathrin (Kukulski et al., 2016). The double mutant strain *swa2-ΔJ sla1Δ* exhibited highly enlarged endocytic coats, which led to highly aberrant actin-driven internalisation of the coat without successful membrane invagination. A stable endocytic coat could be critical in anchoring the actin network to the membrane; this anchoring mechanism could help the actin network in overcoming the opposite force from the cell’s turgor pressure, thus facilitating membrane deformation. Moreover, a precise actin network assembly is crucial to pull both the coat and the membrane underneath. The endocytic actin network primarily polymerises around the rim of the coat, which is where myosin motors also localise (Mund et al., 2018). Controlling its localisation is therefore important, as are the protein ratios to ensure proper anchoring and force generation and transmission throughout the coat. If the coat gets too large, the pulling forces exerted by the actin network are not evenly transmitted to the centre of the coat. In this instance, the effect of turgor pressure cannot be overcome by the actin-driven forces and the coat is detached from the membrane, which remains flat. The higher pulling forces on the edges of the coat compared to its centre would also explain the buckling effect observed during large-coat internalisation events.

Our results show that clathrin coat stabilisation occurs prior to actin network assembly, on a membrane that is, therefore, flat (Kukulski et al., 2012). If the clathrin coat is well ordered, a flat-to-curved transition would be necessary. Without the clathrin triskelia “shuffling” that is thought to be necessary for a flat-to-curved transition to happen, how does the coat accommodate for the rapid final membrane curvature increase? One explanation is the “pre-patterning” of curvature within the coat, which was also postulated in mammalian endocytosis, where gaps between triskelia were observed (Sochacki et al., 2021).

### Conservation of clathrin dynamics

Published FRAP data in mammalian cells show a similar dynamic behaviour for clathrin as we observed in the early phase of endocytosis in yeast (Wu et al., 2001; Loerke et al., 2005; Avinoam et al., 2015). This suggests that the early turnover of clathrin is a highly conserved feature of endocytosis. The late stabilisation has not been so far observed in mammalian cells (Avinoam et al., 2015). However, more work is needed in different cell types and conditions to test whether coat stabilisation occurs during mammalian endocytosis. Many studies point toward mammalian endocytosis being highly variable, depending on cell type, physical and environment cues. Therefore, it is possible that clathrin coat stabilisation might not be a constant feature of mammalian endocytosis, but rather a response to a specific signal. An interesting possibility is that the coat stabilisation could be a mechanism supporting actin-mediated membrane deformation in high turgor pressure or high membrane tension conditions. Studies in mammalian cells have shown that when the membrane tension is high, the actin network becomes essential to deform the membrane (Boulant et al., 2011) and the curvature onset of the clathrin coat can be stalled or delayed (Ferguson et al., 2017), which are two well-established properties of endocytosis in *S. cerevisiae*.

Investigating clathrin dynamics across several organisms and in response to various environment cues would help shed light on a potentially conserved role for coat mechanical properties in response to varying forces required for vesicle formation. This could reveal a crucial mechanism at play in the conserved, yet adaptative nature of the endocytic process.

## METHODS

### Yeast strain generation and maintenance

All yeast strains and plasmids used in this study are listed in **Tables 1 and 2**. All strains were derived from parental strain DDY1102 (*his3-Δ200, leu2-3,112, ura3-52, lys2-801*) and grown on rich glucose medium at 30°C. Fluorescent-protein tagging, gene deletion and C-terminal protein truncations were obtained via homologous recombination with PCR cassettes (Janke et al., 2004), as well as mating and sporulation following standard protocols. *sla1^AAA^*-expressing strain was generated by PCR amplification of the endogenous *SLA1* sequence in two fragments, using primers harbouring the LLD→AAA mutation. Full-length *sla1^AAA^* sequence was cloned into a pFA6a backbone harbouring a hygromycin resistance cassette using Hi-Fi assembly protocol (NEB) and transformed into a seamless-*sla1Δ* strain.

**Table 1:**
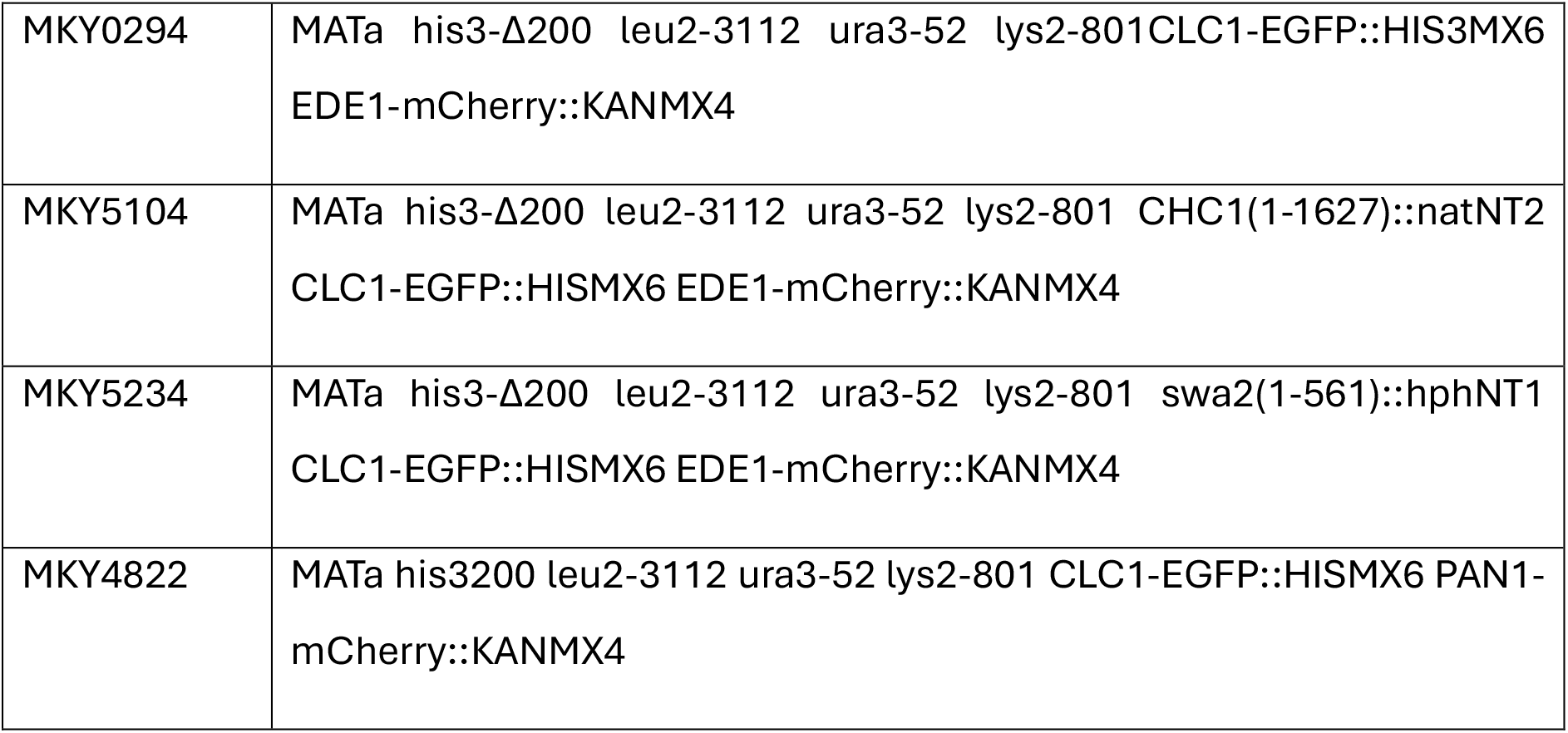
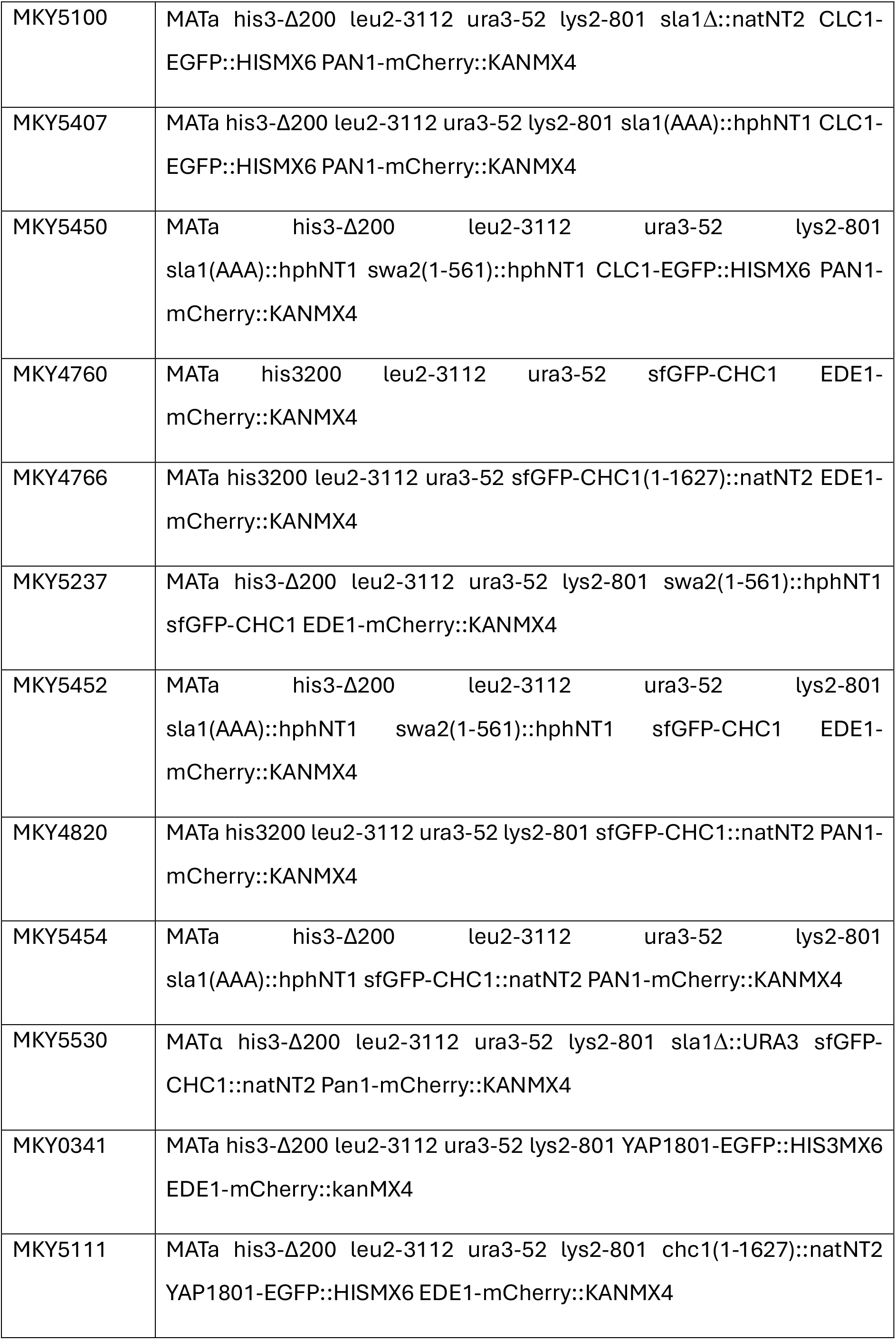
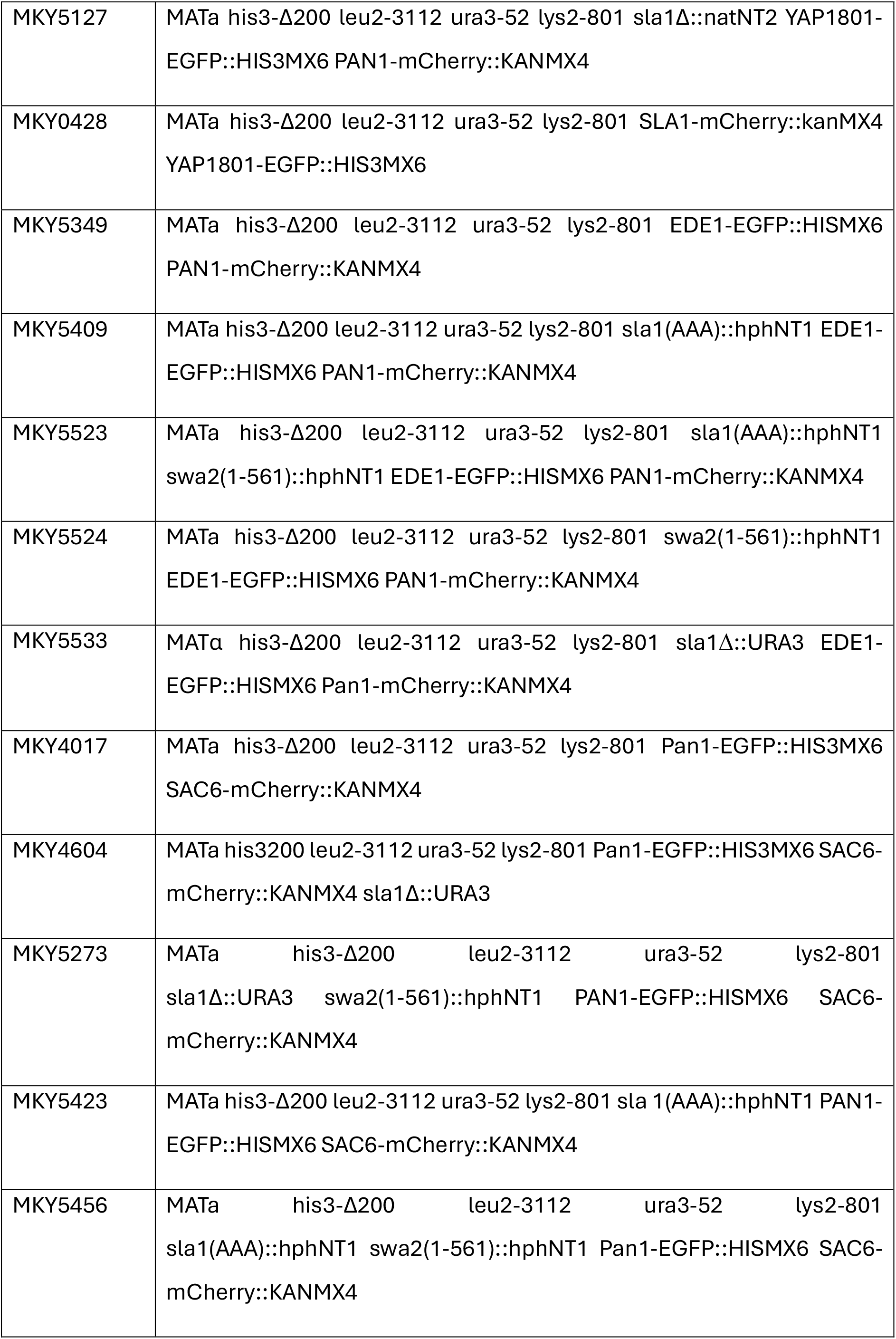
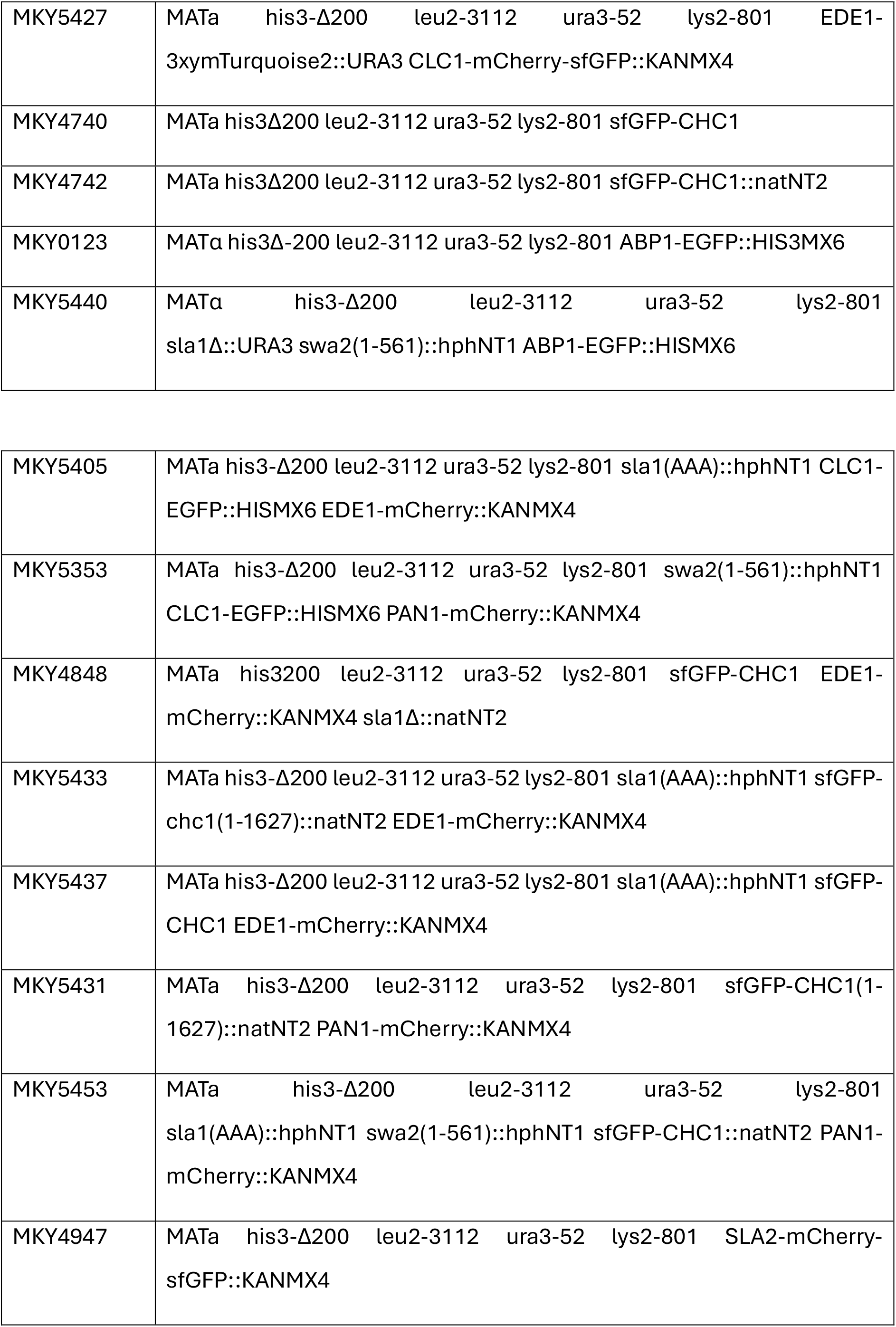
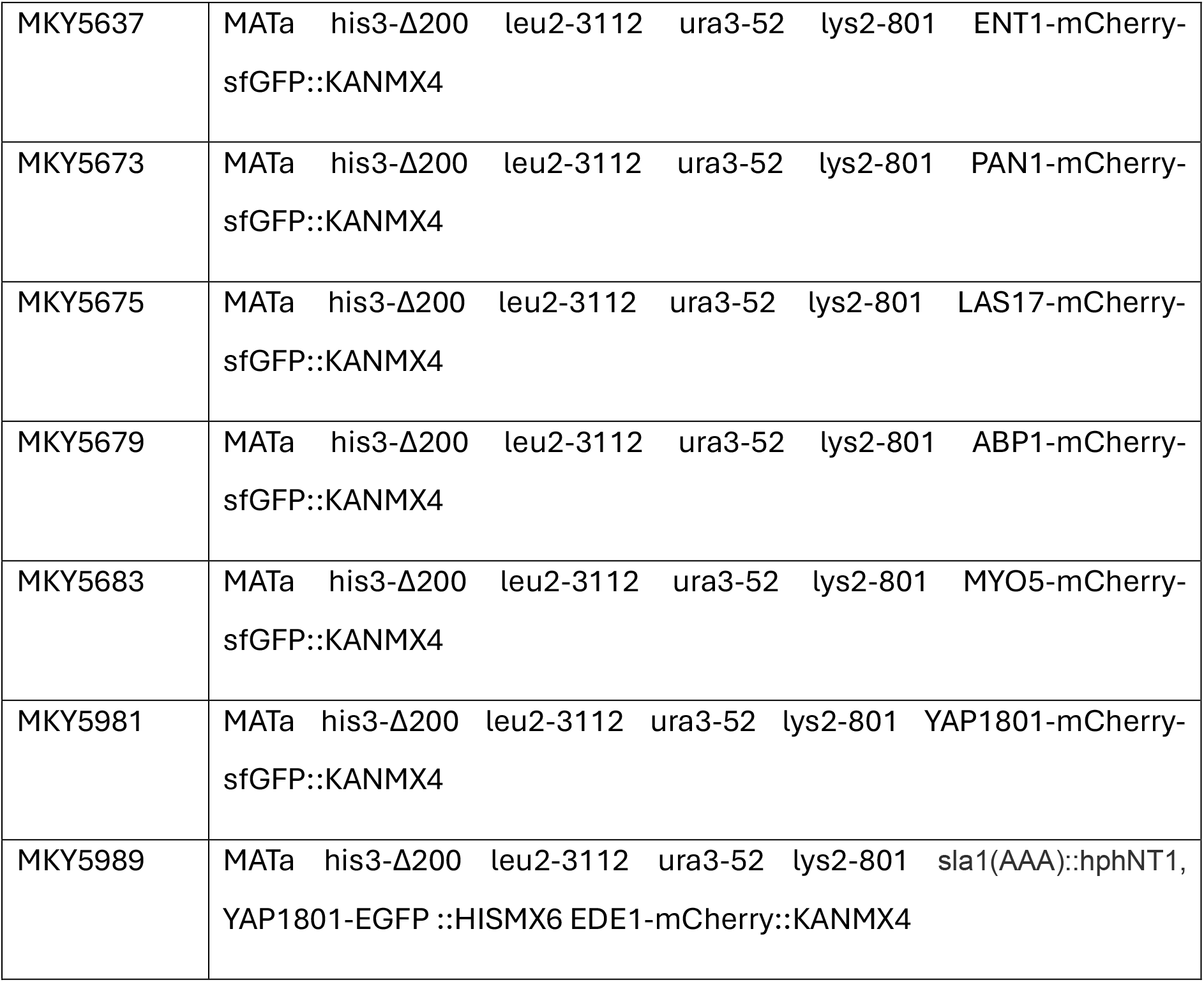
Saccharomyces cerevisiae strains used in this study.

**Table 2:** Plasmids used in this study.

|  |  |
| --- | --- |
| pMK0003 | pFA6a-GFP-HIS3MX6 |
| pMK0005 | pFA6a-mCherry-KanMX4 |
| pMK0019 | pFA6a-natNT2 |
| pMK0052 | pFA6a-hphNT1 |
| pMK0088 | pMaM173 |
| pMK0421 | pMAM17 |
| pMK0421 | pFA6a-mCherry-sfGFP-KanMX4 |
| pMK0660 | pUC57-mCherry-sfGFP-KanMX4 |
| pMK0465 | pSLA1(AAA)-hphNT1 |
| pMK0466 | pSLA1(AAA)-EGFP-hphNT1 |
| pMK0099 | pFA6a-EGFP-natNT2 |

### Live fluorescence microscopy imaging

#### Sample preparation

Yeast cells were grown to OD_600_ 0.3-0.8 in SC-Trp at 24°C. The cells were then adhered onto a Concanavalin-A-coated coverslip and washed with fresh medium immediately prior to imaging.

#### Widefield epiffuorescence imaging

Images were acquired by an an ORCA ER-CCD camera (Hamamatsu) using an Olympus IX-81 microscope equipped with a 100x/NA1.45 oil objective. For single-colour acquisition, Sapphire 488-100 (488nm) or Compass 315M (561nm) lasers (Coherent) were used as the illumination source. For 2-colour imaging, an XCITE 120PC metal halide lamp (EXFO) was used in combination with U-MGFPHQ and U-MRFPHQ filter sets (Olympus) for excitation and emission light filtering.

#### Total Internal Reffection Fluorescence (TIRF) microscopy

Timelapses were acquired by an ImageEM X2 EM-CCD camera (Hamamatsu) using a widefield Olympus IX-83 microscope equipped with a 150x/NA1.45 oil objective. 3xymTurquoise2-, EGFP/sfGFP- and mCherry-tagged proteins were illuminated with 405nm, 488nm and 561nm lasers, respectively. Excitation and emission lights were filtered using a TRF89902 405/488/561/647 nm quad-band filter set (Chroma).

Microscope and laser angles were controlled by the VisiView software (Visitron) and iLas2 system (Roper Scientific), respectively. For dual-colour imaging, 488nm and 561nm lasers were used simultaneously for sample illumination. The emission lights were then split using a Gemini system (Hamamatsu) equipped with emission filters FF03-525/50-25 and FF01-630/92-25, in combination with a dichroic mirror Di03-R488/561-t1−25 × 36 (Semrock Brightline). Chromatic aberration correction was performed using 1:100 TetraSpeck beads in SC-Trp.

#### Fluorescence Recovery After Photobleaching (FRAP)

Photobleaching of EGFP/sfGFP and mCherry was performed using 488nm and 561nm lasers, respectively. Spatial targeting of endocytic patches was achieved with the iLas2 targeting system.

### ATP depletion

Yeast samples were prepared for imaging according to standard protocols (see **section 6.a.**). ATP was depleted from yeast cells 30min prior to imaging by incubating the cells in Synthetic Complete medium without Tryptophane (SC-Trp) supplemented with 20mM 2- deoxyglucose and 0.2mM sodium azide (NaN_3_). The cells were then imaged within a 30- min timeframe.

### Correlative Light and Electron Microscopy (CLEM)

For CLEM, sample preparation, FM, ET and correlation was performed as described previously (Kukulski et. al, 2011). Briefly, *S.cerevisiae* strains expressing Abp1-eGFP in *swa2-ΔJ sla1Δ* and WT backgrounds were grown in low fluorescence medium at 24°C until they reach the log phase. The cells were pelleted using vaccum filtaration and cryo fixed by high pressure freezing done with Leica EM ICE high pressure freezer.

Further infiltration was done by freeze substitution using Leica AFS2. FS occurred at −90°C for 48–60 h with 0.1% (wt/vol) uranyl acetate in glass-distilled acetone. The temperature was then raised to −45°C (5°C/hour), and samples were washed with acetone and infiltrated with increasing concentrations (10, 25, 50, and 75%; 4 h each) of Lowicryl in acetone while the temperature was further raised to −25°C. 100% Lowicryl was exchanged three times in 10-h steps and samples were UV polymerized at −25°C for 48 h, after which the temperature was raised to 20°C (5°C/hour) and UV polymerization continued for 48 h. 280-nm sections were cut with a microtome and a diamond knife and picked up on carbon-coated 200 mesh copper grids.

Followed by FM on olympus IX81, dual-axis tilt series were collected at a pixel size of 3.2916 nm and at 1 degree increments using Talos L120C. The IMOD software package was used for tomogram reconstruction. Correlation of the fluorescent spots with coordinates of corresponding electron tomograms was performed by using fluorescent microspheres as a fiducial system.

2 and 3 independent tomograms were acquired and analysed for wild type and *swa2-ΔJ sla1Δ* cells, respectively.

### Data analysis

#### FRAP analysis

All movies were corrected for photobleaching and cytoplasmic background using the “subtract background” FiJi plugin with a 5-px rolling ball radius, and calculating the ratio between the background-corrected image stack and the corresponding median-filtered image stack (spot radius 50px).

#### 2-colour FRAP

For average FRAP analyses, individual photobleaching events were aligned to their time of photobleaching (t=0) and normalised using the average mCherry fluorescence intensity of five seconds prior to photobleaching as the maximum, and the fluorescence intensity immediately after photobleaching as the minimum. The GFP signal was normalised to the mCherry curve by affine-function fitting of each individual GFP curve to its corresponding mCherry signal from a 6-second window prior to photobleaching.

For single event FRAP analysis, the GFP fluorescence intensity curves were smoothened and normalised and used for the subsequent normalisation of the mCherry fluorescence signal. The fluorescence recovery after photobleaching (marked as t=0) for single events was smoothened and an exponential plateau model was fitted to the data, in order to extract the characteristic parameters Y_0_, Y_m_ and t_1/2_ (respectively: fluorescence intensity at t=0; calculated maximum of fluorescence reached post photobleaching; and half-time of the fluorescence recovery). The minimum and maximum of the GFP signal was also extracted from the smoothed curve, and the endocytic event endpoint was identified by extracting the time point for which the curve reached a low intensity threshold (i.e. average fluorescence signal recorded in absence of any endocytic event).

To identify significant fluorescence recoveries, we set a 0.2 threshold to distinguish fluorescence recovery from lack thereof, accounting for the high noise associated with fluorescence curves from single events.

#### Single-colour FRAP

Individual photobleaching events were aligned to their time of photobleaching (t=0) and normalised using the average fluorescence intensity of ten seconds prior to photobleaching as the maximum, and the fluorescence intensity immediately after photobleaching as the minimum. To extract the characteristic parameters Y_0_, Y_m_ and t_1/2_, an exponential plateau model was fitted to the mean fluorescence recovery. Mean fluorescence intensity and its associated standard error were plotted.

#### Endocytic protein lifetime quantification

The protein lifetime was measured from endocytic events imaged by TIRF microscopy. All movies were corrected for photobleaching and cytoplasmic background (see previous section). The fluorescence intensities over time were extracted from the corrected movies; start and endpoints of the endocytic events were identified by extracting the time point for which the curve reached a low intensity threshold (i.e. average fluorescence signal recorded in absence of any endocytic event).

#### Endocytic protein ffuorescence intensity quantification

For long-lived proteins displaying a fluorescence intensity plateau (i.e. Ede1 and Clathrin), the fluorescence intensity was calculated from a median intensity projection over 15 seconds. The fluorescence intensity of late-phase proteins was estimated by calculating the mean fluorescence intensity of five frames centred around the fluorescence maximum.

#### Inward movement determination

All movies were corrected for photobleaching and cytoplasmic background using a FiJi plugin developed in our lab (Picco and Kaksonen, 2017). The Particle Tracking plugin from the Mosaic suite on FiJi was used to track the movement of Pan1-GFP patches. These movements were then aligned using an algorithm developed in our lab (Picco and Kaksonen, 2017).

## Statistical analysis

Statistical tests were performed using the GraphPad Prism software.

Yeast strains and plasmids

## Acknowledgements

We thank all members of the Kaksonen and Roux labs for their insights and feedback. We also extend our warmest thanks to Karsten Kruse for the fruitful discussions and insights about our 2-colour FRAP setup, Andrea Picco for his help with image analysis, and Sofia Meghezi for her assistance. We are also grateful for the kind feedback from Liz Miller, Leonardo Almeida-Souza and Robbie Loewith. This work was supported by the Swiss National Science Foundation grants to M.K. (#310030_212288) and to A.R. (#CRSII5_189996 and #310030_200793) and the European Research Council Synergy grant to A.R. (#951324-R2-TENSION).

