## Supplemental figures for "A switch in clathrin turnover controls endocytic coat size and organisation"

#### Supplementary figures

### Figure S1

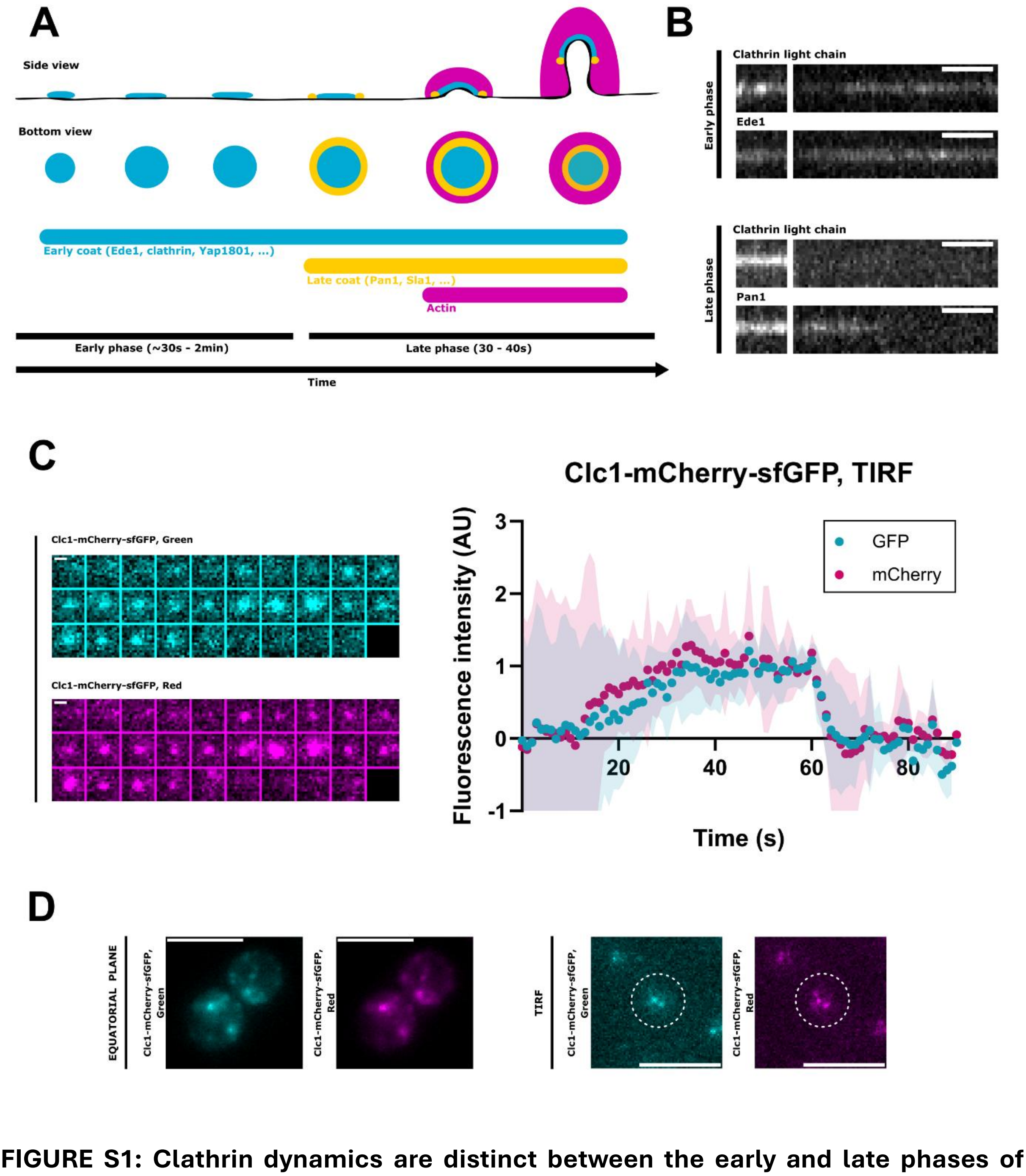


**FIGURE S1: Clathrin dynamics are distinct between the early and late phases of endocytic site maturation. A.** Schematic representation of endocytic site maturation in yeast. **B.** Representative image of clathrin-GFP FRAP during the early phase (top) and the late phase (bottom, using Ede1-mCherry and Pan1-mCherry as phase markers (respectively). Scale bar: 5 seconds. **C.** Left: Representative example of a timelapse of clathrin-mCherry-sfGFP recruitment at endocytic sites, imaged with simultaneous 2-colour TIRF. Framerate: X seconds. Scalebar: Xnm. Right: average clathrin-mCherry-sfGFP recruitment at endocytic sites. Error bars represent SEM. N=16. **D.** Left panels: Widefield imaging of the equatorial plane of a yeast cell expressing Clc1-mCherry-sfGFP endogenously. Right panels: TIRF imaging of of a yeast cell expressing Clc1-mCherry-sfGFP endogenously. Dotted circles represent the cell outline. Scale bars: 4μm.

### Figure S2

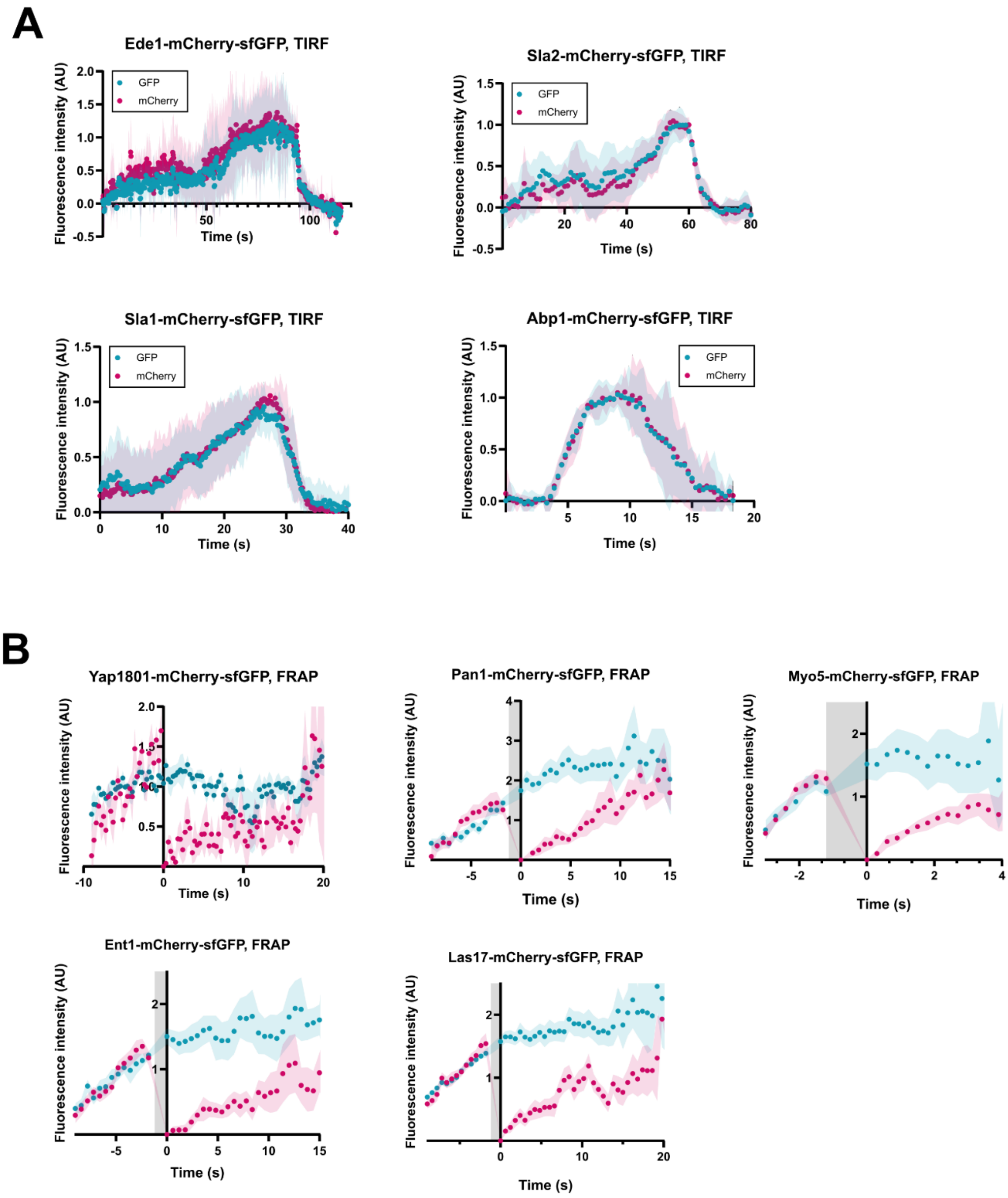


**FIGURE S2: Endocytic coat rigidifies over time. A.** Average Ede1, Sla2, Sla1 and Abp1 recruitment at endocytic sites All proteins were expressed as fusion proteins with tandem mCherry-sfGFP, and imaged with simultaneous 2-colour TIRF. For all proteins, N=16. Error bars represent standard deviation. **B.** Average mCherry FRAP compared to GFP fluorescence intensity, for early-coat protein Yap1801 (N=14), mid-coat protein Ent1 (N=47), late-coat proteins Pan1 (N=48) and Las17 (N=25), and actin-module protein Myo5 (N=61). Gray areas represent photobleaching durations. Error bars represent standard errors associated with the mean (SEM).

### Figure S3

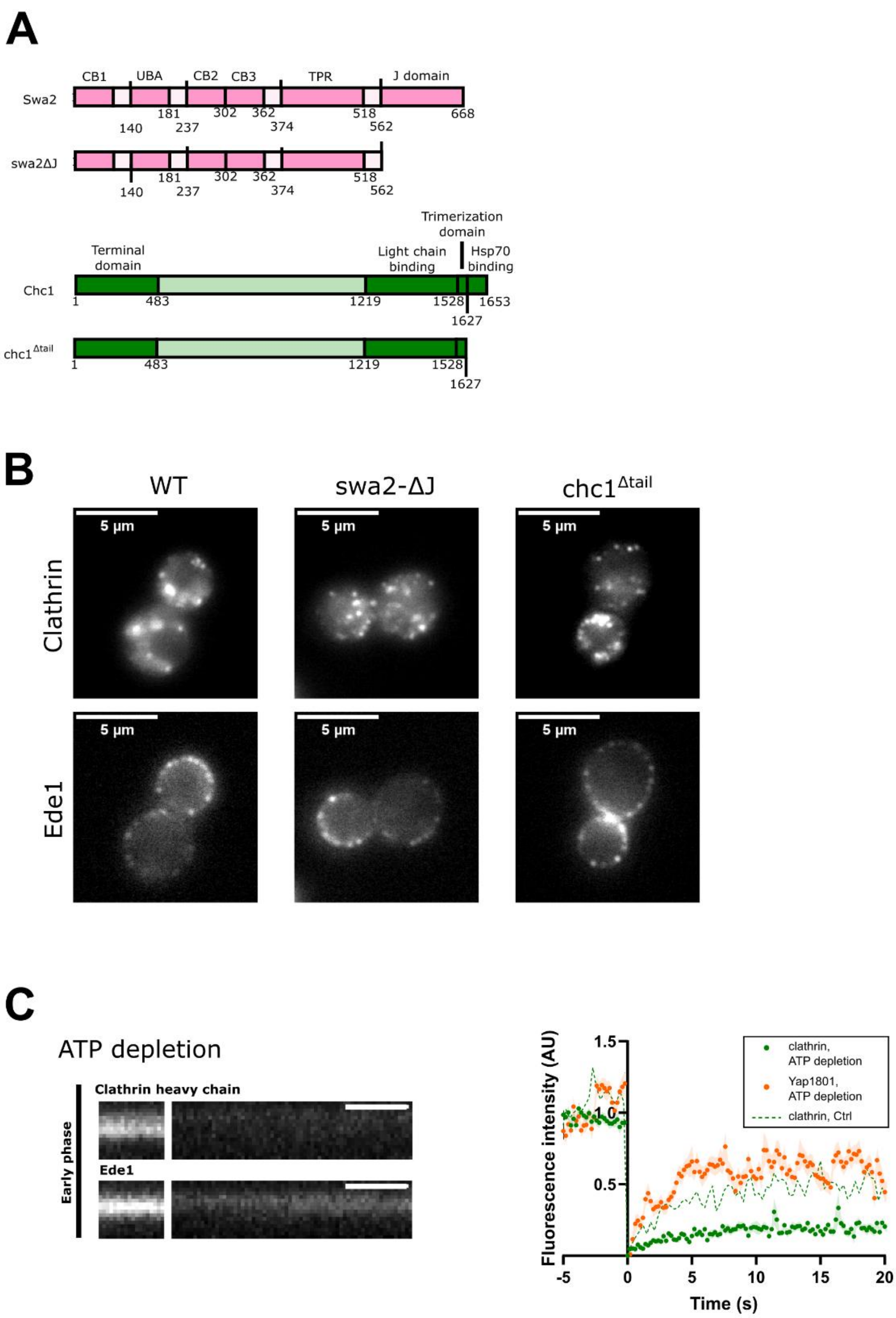


**FIGURE S3: Early-phase clathrin turnover depends on Swa2/ATPase. A.** Overview of the sequence domain for WT Swa2 and clathrin heavy chain, and for clathrin-uncoating mutants swa2-ΔJ, lacking the J domain recruiting the ATPase, and chc1$^{\Delta tail}$, lacking the unstructured C-ter tail binding to the ATPase. **B.** Widefield microscopy imaging at the equatorial plane of strains expressing sfGFP-Chc1 (top) and Ede1-mCherry (bottom) in wild type and Swa2/ATPase-abolishing mutant backgrounds. **C.** Fluorescence recovery after photobleaching of Clc1-GFP and Yap1801-GFP during the early phase of ATP-depleted cells. Left: representative kymograph of ATP-depleted cells expressing GFP-Chc1 and Ede1-mCherry. Scale bar: 5 seconds. Right: Mean FRAP for Clathrin and Yap1801 in ATP depleted cells. Clathrin: N=23; Yap1801: N=60. Error bars represent SEM.

### Figure S4

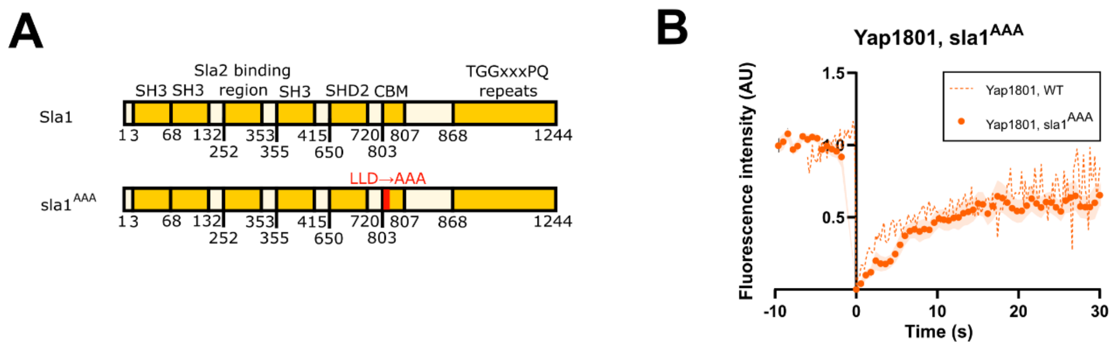


**FIGURE S4: Clathrin adaptor Yap1801 is unaffected by alterations in late-phase clathrin coat dynamics. A.** Schematic representation of the known domains in the Sla1 sequence (top) and localisation of the mutation harboured by the sla1$^{AAA}$ mutant (bottom). **B.** Mean fluorescence recovery after photobleaching of Yap1801-GFP during the late phase of yeast cells expressing point mutant sla1$^{AAA}$ (N=24), compared to that observed in WT cells (N= 35). Error bars represent SEM.

### Figure S5

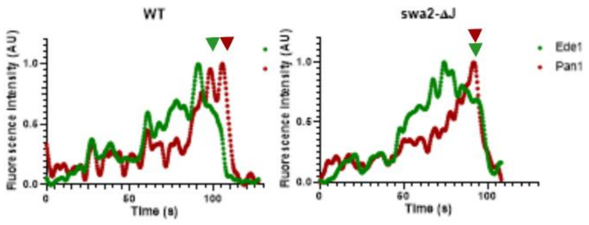


**FIGURE S5: Impaired early-phase clathrin turnover hampers Ede1 disassembly from endocytic sites.** Fluorescence intensity over time of Ede1-GFP and Pan1-mCherry in a single endocytic event from wild type cells (left) and swa2-ΔJ mutants (right). Arrows highlight the fluorescence decrease onset for Ede1 (green) and Pan1 (red).

### Figure S6

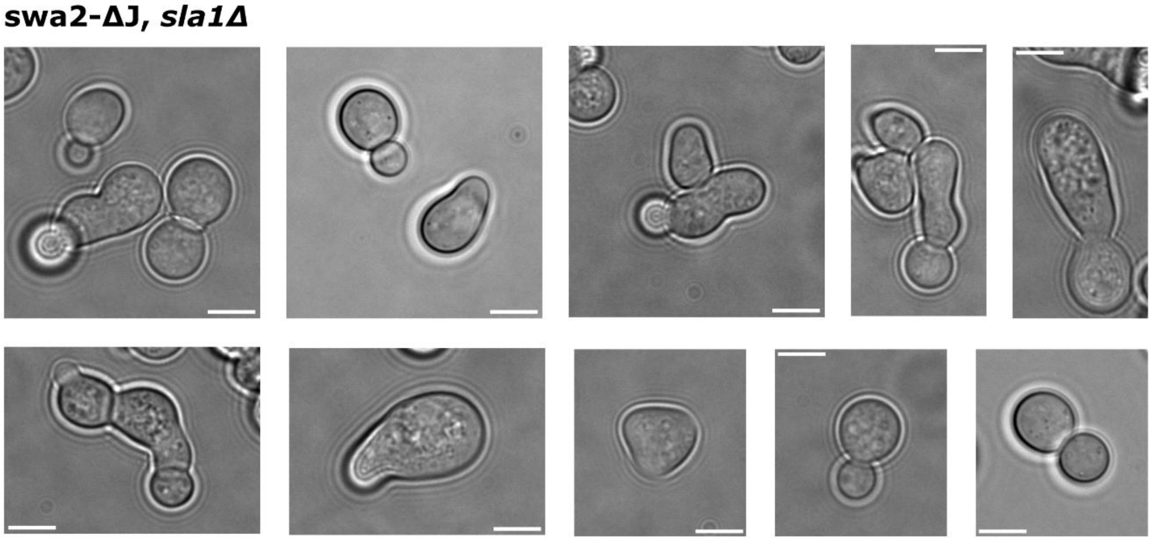


**FIGURE S6: Impaired control of clathrin assembly at endocytic sites decreases cell fitness.** Brightfield images of swa2-ΔJ *sla1Δ* mutants. Scale bar: 5μm.
